# Distributed evidence accumulation across macaque large-scale neocortical networks during perceptual decision making

**DOI:** 10.1101/2023.12.26.573347

**Authors:** Licheng Zou, Nicola Palomero-Gallagher, Douglas Zhou, Songting Li, Jorge F. Mejias

**Affiliations:** Cognitive and Systems Neuroscience Group, Swammerdam Institute of Life Sciences, University of Amsterdam, Netherlands; Zhiyuan College, Shanghai Jiao Tong University, China; Institute of Neuroscience and Medicine (INM-1), Research Centre Julich, Germany; C. & O. Vogt Institute of Brain Research, Heinrich-Heine-University Düsseldorf, Germany; School of Mathematical Sciences, MOE-LSC, Institute of Natural Sciences, Shanghai Jiao Tong University, China

**Author notes:** Corresponding authors: Songting Li, Jorge F. Mejias.

## Abstract

Despite the traditional view of parietal cortex as an important region for perceptual decision-making, recent evidence suggests that sensory accumulation occurs simultaneously across many cortical regions. We explored this hypothesis by integrating connectivity, cellular and receptor density datasets and building a large-scale macaque cortical model able to integrate conflicting sensory signals and perform a decision-making task. Our results reveal sensory evidence accumulation supported by a distributed network of temporal, parietal and frontal regions, with flexible sequential bottom-up or top-down modulation pathways depending on task difficulty. The model replicates experimental lesioning effects and reveals that the causal irrelevance of parietal areas like LIP for decision performance is explained by compensatory mechanisms within a distributed integration process. The model also reproduces observed temporal gating effects of distractor timing during and after the integration process. Overall, our work hints at perceptual integration during decision-making as a broad distributed phenomenon, providing multiple testable predictions.

## Introduction

Decision-making (DM) is a fundamental cognitive function in the brain, involving the processing of sensory information towards a categorical decision and a subsequent motion response (*1*). Electrophysiological recordings have characterized DM as an evidence accumulation process, where a decision is made when the firing rate of specific neuronal populations, typically in association parietal areas such as LIP, exceeds a threshold (*2, 3*). This principle of sensory integration within local (parietal) circuits has been used to understand a vast range of decision-making paradigms, from simple perceptual decision-making tasks (*4*–*7*) to context-dependent choices (*8*–*10*), multisensory integration (*11*–*14*) or even perceptual choices in the context of consciousness or arousal modulation (*15*–*18*).

In recent years, however, compelling evidence has emerged against the local (parietal) sensory integration theory. Notably, lesioning and optogenetic studies in rodents have revealed that the inactivation of parietal cortex has little effect on the performance of decision tasks, even though the activity of those areas is highly correlated with decision speed and accuracy (*19, 20*). Similar experiments in macaques led to similar conclusions, suggesting that sensory areas like MT have a greater influence on achieving good performance than LIP (*21*). It seems implausible, therefore, that sensory integration necessary to guide decisions would occur solely on parietal circuits. Recent studies have explored whether the involvement of multiple brain regions could support perceptual decision-making (*9, 22*–*24*), revealing a wide range of differentiated roles across cortical regions (*25, 26*) and emergent interactions across different areas (*9*). Promising conceptual advances have been facilitated by computational modeling work involving two or three interacting areas (*27*–*29*). However, the lack of a data-constrained, brain-wide framework to compare and evaluate the contributions of individual areas involved in decision-making tasks has hindered further progress.

In this work, we integrated multiple brain-wide data sets and used them to develop a computational model of the large-scale network of the macaque neocortex with the capacity to simulate decision-making tasks. The data integrated into the model included cortex-wide variations in autoradiography-derived NMDA and GABA_A_ receptor densities per neuron, and average dendritic spine count per cell (*30*–*32*), which contribute to capture part of the existing regional heterogeneity in cortex. By replicating a classical perceptual decision-making task (random dot motion discrimination), our model showed the involvement of multiple cortical areas in sensory accumulation, in contrast with the traditionally assumed local integration in parietal cortex. We also observed different roles, such as sensory, accumulators and categorizers, emerging across cortical areas. In addition, the model revealed the existence of two distinct dynamical regimes governing the flow of information during sensory integration, with the level of stimulus coherence determining which of them drove the decision process. Furthermore, we proposed that temporal gating during decision-making, a phenomenon observed in mice so far, would be present in macaques depending on the robustness of the distributed network supporting integration and maintenance of choice memory (*33*). Lastly, by simulating lesions in different cortical regions, we reproduced the results of experimental inactivation studies, supporting the hypothesis of the causal irrelevance of area LIP for decision making and proposing a more important role for temporal lobe regions.

## Results

### A large-scale macaque cortex model incorporating data from multiple macroscopic gradients

We combined multiple neuroanatomical data sets with local dynamical models to build a large-scale model of the macaque cortex (Figure 1A; see Materials and Methods for further details). We first considered a local circuit of two stimulus-selective excitatory populations and one non-selective inhibitory population. This local circuit was our template to describe each of the 40 cortical areas considered in our large-scale model, with properties of each individual circuit varying across areas following available area-specific data. This ensures that each cortical area was unique in its configuration and dynamics, and reflected the high level of neural and circuit heterogeneity found in real brains. The area-specific properties revealed macroscopic gradients across cortex (*34*), and included variations in the effective strength of synapses inferred from local properties such as the number of dendritic spines per pyramidal cell (*31, 33*) and the area-specific density of NMDA and GABA_A_ receptors per neuron (*32, 35*)(Figure 1B).

**Figure 1:**
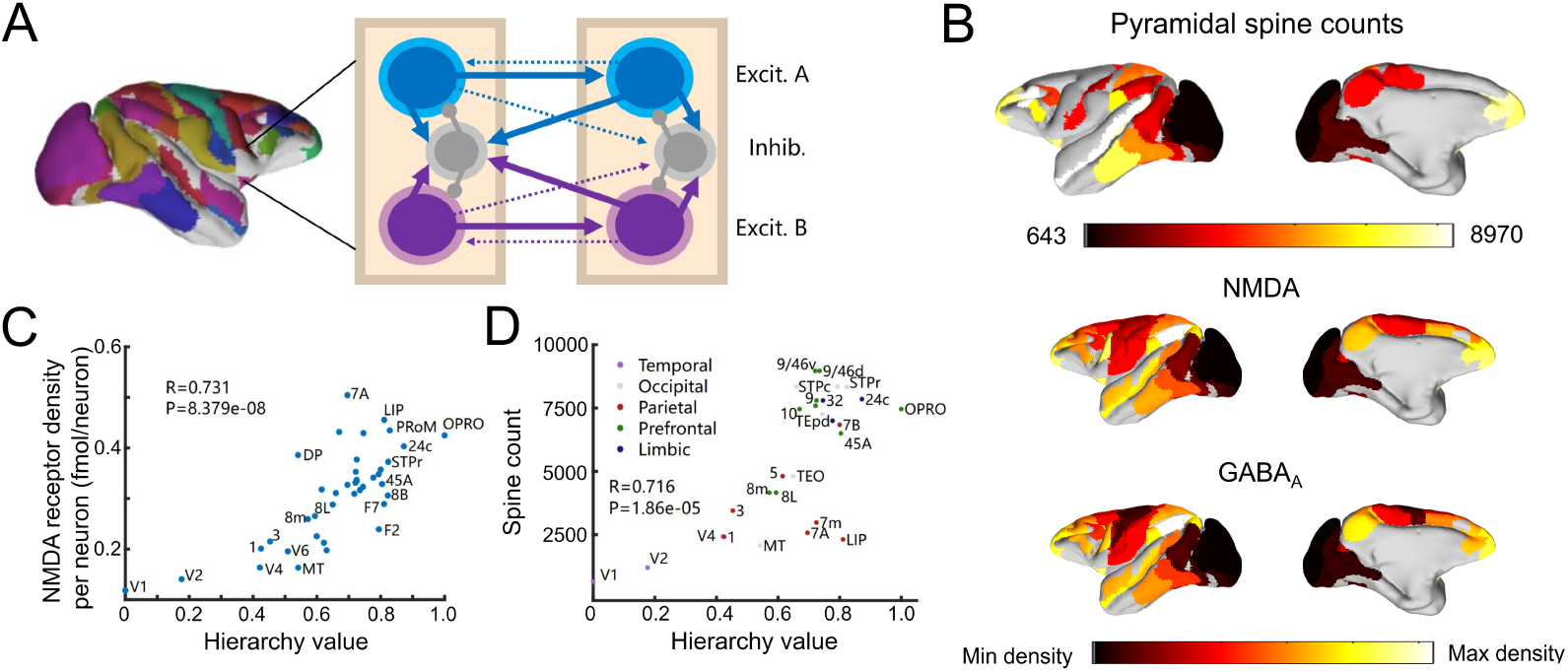
Scheme and anatomical basis of the large-scale macaque cortical model. (A) Lateral view of the macaque cortical surface with modelled areas in color. In each area, cortical dynamics follow a local winner-take-all model, and areas are connected using anatomical connectivity data. (B) Area-specific values for the number of dendritic spines per neuron (top), the densities of NMDA receptors per neuron (middle) and GABA_A_ receptors per neuron (bottom). (C) Correlation between NMDA receptor density per neuron and anatomical hierarchy as defined by layer-dependent connections. (D) Correlation between spine count data and anatomical hierarchy (labels of areas were partly displayed for better visualization).

Before assembling these areas into a large-scale cortical model, we analyzed the organization of the receptor density per neuron data (*32*) when compared to the position of each area in the anatomical hierarchy (*36, 37*) and area-specific dendritic spine numbers (*31, 33*). A strong positive correlation was found between NMDA receptor density and the cortical hierarchical position inferred from laminar connectivity data (r=0.731) (Figure 1C), and NMDA receptor density had an almost linear relationship with GABA receptor density (Suppl. Fig. S1). As previously shown (*33, 38*), dendritic spine count positively correlated with cortical hierarchy (Figure 1D). We identified here some parietal areas (such as LIP, 7m, and 7a) as outliers based on their low pyramidal spine count but high cortical hierarchy value and NMDA receptor density. This suggested that NMDA-mediated synapses in those areas preferentially target inhibitory neurons.

The data-constrained local circuits were then connected between them to form an extended cortical network, using tract-tracing connectivity data of the macaque neocortex (*36, 37, 39, 40*), leading to a cortical model constrained at both the local and large-scale levels by neuroanatomical data. Using the layer-specificity and projection-directionality provided by this dataset, and following previous work (*33, 41*), we implemented a counterstream inhibitory bias in our network, which assumes that feedforward and feedback projections along the cortical hierarchy slightly but preferentially target excitatory and inhibitory neurons, respectively. Background input to cortical areas was varied across areas to mimic the differentiated thalamocortical projections across cortex and to facilitate that all cortical areas displayed a similar spontaneous activity level of about 0.5Hz.

### Distributed and heterogeneous evidence accumulation process in decision making

We simulated our large-scale macaque cortical model performing a classical two-choice random-dot motion discrimination task for decision making (*2, 3*), where subjects have to choose which of two possible movement directions has a larger number of dots involved (Figure 2A). In our model, this visual input is modelled as two 700-ms-duration external currents entering both selective excitatory populations in V1 (Figure 2A and B), with dot movement coherence (the degree of agreement among moving dots, ranging from 0-100%), or motion contrast (the consistency in the direction of motion for all moving dots, ranging from -1 to 1), reflected in the difference between these currents (Figure 2B). This sensory input drives a competition between both selective excitatory V1 populations, and subsequently such a competition took place in other cortical areas within individual excitatory populations.

**Figure 2:**
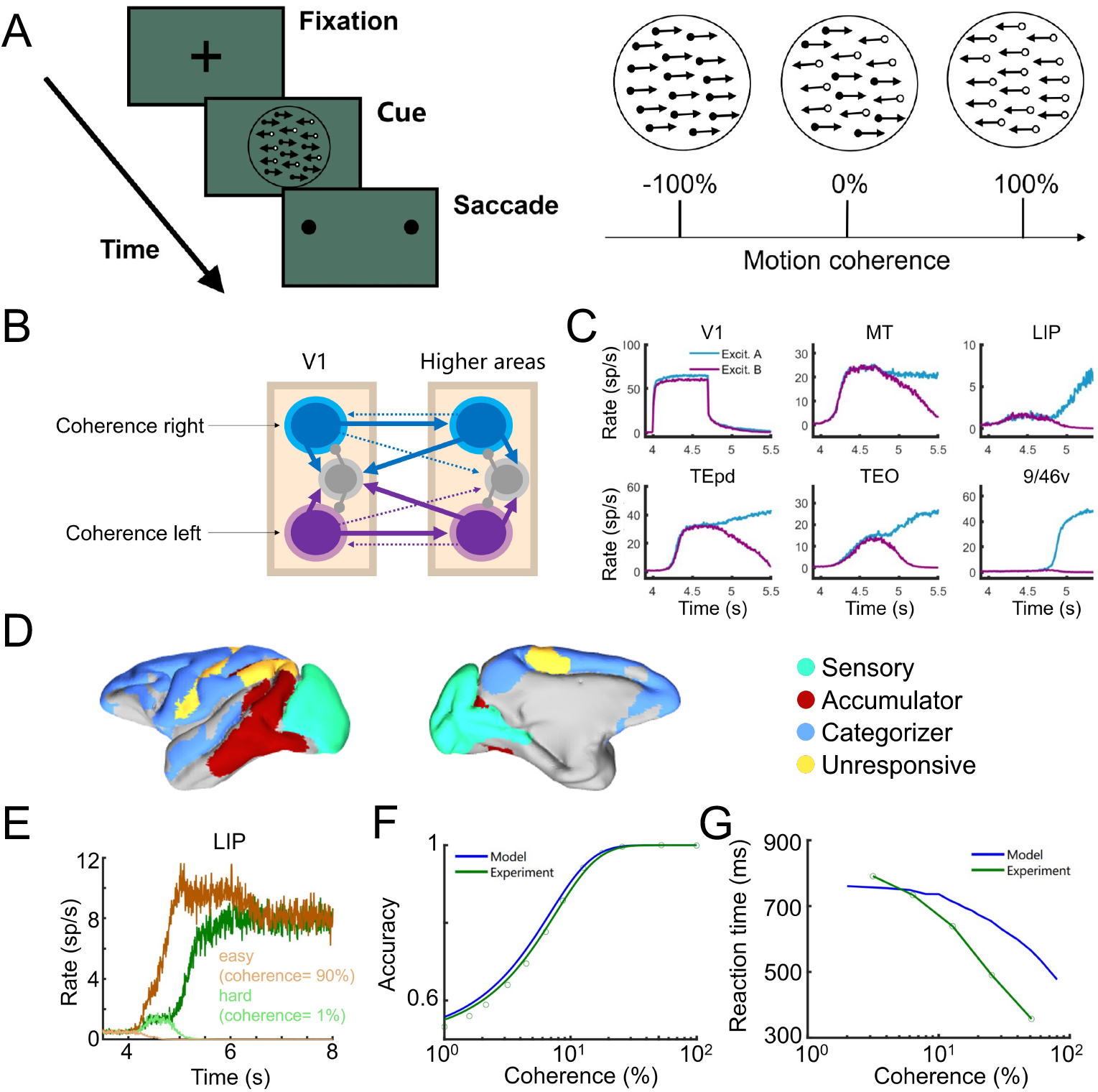
Large-scale cortical model reveals a distributed sensory accumulation process for decision making. (A) Random-dot motion discrimination task simulated in our work (left panel), where monkeys must choose which is the preferred direction of movement of the visual stimuli. The stimuli can display any coherence level from 100% right to 100% left movement (right panel). (B) Circuit schematic of the simulation. Two external stimuli representing motion information enter the excitatory populations of V1. (C) Activity of selected cortical areas during the decision-making task, with a 5%-coherent selective visual input duration of 700 ms, displaying a sensory accumulation process distributed across areas. (D) Clusters of all 40 areas according to characteristic responses. (E) Sensory accumulation activity in LIP, for two different coherence levels. Green (orange) traces correspond to hard (easy) trials. Populations encoding the correct and incorrect choice are shown in thick and thin lines, respectively. (F) Psychometric and (G) chronometric curves as a function of stimulus coherence. Green dots and curves are experimental data from macaques **(*2*)**.

Simulations of the large-scale cortical model (Figure 2C) revealed that the competing visual stimuli induced the accumulation of sensory evidence and the resulting competition and winner-take-all dynamics in area LIP, as shown in experiments and classical models (*6, 7*). However, such dynamics were also present in other brain areas, such as temporal lobe regions (TEO, TEpd), prefrontal cortex (9/46v), and many others (Suppl. Fig. S2). This suggests that the decision process, including sensory accumulation and winner-take-all dynamics, occurs across a distributed cortical network rather than in localized parietal regions. When comparing the firing rate evolution of stimulus-selective sustained activity across 40 areas and clustered them according to characteristic dynamics (Figure 2D), we found dynamics greatly varied across the cortex: sensory areas like V1 and V2 responded during cue duration and maintained spontaneous states during delay (sensory), somatosensory areas like 1 and 3 displayed little reaction to the stimulus (unresponsive), frontal areas like F5 and 9/46v underwent a sharp transition reflecting a categorical choice dynamic (categorizer), areas like LIP, TEO and 7M showed a slow and gradual ramping process, and temporal areas like MT and TEpd displayed mutual ramping after stimulus onset and then bifurcated to encode the final decision (accumulator). As in classical models and experimental evidence on single areas, the accumulation process was dependent on the level of stimulus coherence (Figure 2E): as illustrated by this example in LIP, ramping activities bifurcated earlier in high-coherence trials compared to low-coherence ones, and experienced different activity patterns for two situations. In both cases, the winning population displayed sustained activity after the integration process, due to the bistability of the system, and related to working memory capacity of the network (*33*). When adopting a majority rule to infer the behavioral decision from the winning populations across all cortical areas, the large-scale cortical model was also able to replicate psychometric curves in agreement with macaque behavioral output data (Figure 2F), as well as plausible chronometric curves (Figure 2G). The reaction time in chronometric curves was defined as the time at which mean firing rates of excitatory populations displaying bifurcation (like LIP, 9/46v) reached a threshold of 3 Hz to fit experimental data, signaling a joint contribution of association areas towards a decision.

### Winning onset times uncover motion information flow across the cortex

While evidence accumulation was observed across many cortical areas, integration times greatly varied across regions. We analyzed the temporal integration across multiple areas by comparing the normalized firing rate traces of the populations encoding the winning choice. For low coherence levels, temporal and prefrontal areas (like TEO and 9/46v, respectively) led the integration and were the first ones to reach their half-maximum level (Fig. 3A, left). However, the situation changed for trials with high coherence levels, as visual areas like MT were the first ones reaching their half-maximum level (Fig. 3A, right) and temporal and frontal areas evolved at a slower pace. This suggests that the information flow during decision-making is stimulus-dependent, and that the functional relationships between cortical areas (i.e. which area leads the integration), rather than being a fixed structure, flexibly depends on task difficulty.

**Figure 3:**
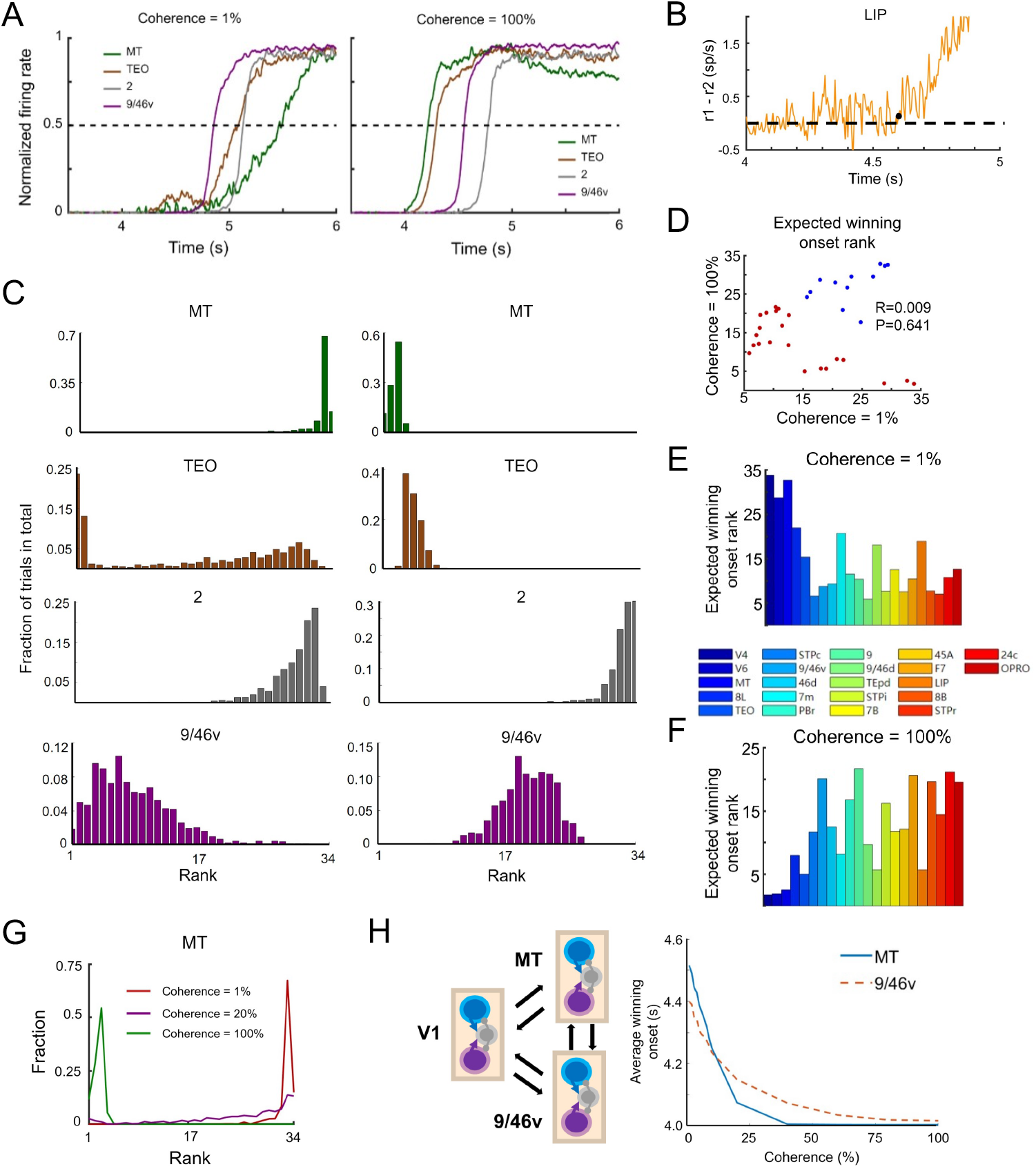
Winning onset analysis reveals information flow during decision making. (A) Normalized firing rate difference between both excitatory populations for selected cortical areas under zero (left) and full coherence (right). (B) Winning onset was defined as the time from stimulus onset until the difference of firing rate between both excitatory populations rose from zero (black dot), without reverting to zero later in the trial. Dotted line indicates the baseline for excitatory firing rate difference. (C) Winning-onset rank distribution of selected areas under zero (left) and full coherence (right). (D) Expected (average) winning-onset rank for association areas during low coherence (1%) versus high coherence (100%) trials. Red dots corresponded to areas exhibiting negative correlations in both scenarios, and blue dots were areas remaining stable. (E) Expected winning-onset rank tended to decrease along hierarchy during low-coherence situation. (F) Expected winning-onset rank tended to increase along hierarchy during high-coherence situation. Selected areas in (E) and (F) corresponded to red points in (D). (G) Winning-onset rank for area MT across six coherence levels. (H) Toy distributed model consisting of three areas (left) and the relation between coherence level and winning onset for MT and 9/46v. All datapoints in (C) -(H) were averaged over 2000 trials.

To estimate the moment in which a winner emerges in the local competition between selective populations in each area, we defined the ‘winning onset’ in a cortical area as the moment in which the difference between their firing rates becomes larger than a certain threshold (zero, for simplicity) and continues evolving towards the decision threshold without returning back (black dot in Fig. 3B). Intuitively, the winning onset signals the time at which a winner population gains an edge over the other one, and a differentiated integration emerges. The winning onset provides a simple way to evaluate which areas lead the integration process, which aligns reasonably well with decision sequences observed in categorical decision tasks in macaques (*9*)(see Suppl. Fig. S3 for a direct comparison). We simulated, for a given coherence level, 2000 decision-making trials and computed the winning onset for each area. By sorting these winning onset times and ranking all areas from first/fastest to last/slower, we obtained a trial-averaged distribution of winning-onset ranks for each area (Fig. 3C).

For low coherence values, frontal areas like 9/46v and 45A, as well as temporal areas like TEO, STPc and STPi were usually ranked in the first place. Areas linking sensory and association areas, like MT and LIP, as well as vision-unrelated areas such as areas 2 and 5 (primary and secondary somatosensory cortex), fell behind in the ranking. This suggests that, under low coherence input, higher association areas led the sensory integration and decision process, which was later reflected in the activity of areas like LIP and MT. For high coherence values, the pathway for sensory integration drastically changed (Fig. 3C, right). In this condition, LIP, MT and similar areas connecting sensory and association cortex become fast integrators and reclaim the first places in the winning onset ranking. Association areas, on the other hand, became slower and decreased in the ranking (although some of them, like TEO, remain in moderately high ranks). Overall, high-ranked areas seemed to be hierarchically close to early sensory areas V1 and V2, indicating that sensory integration follows the anatomical hierarchy for high coherence input.

Two distinct mechanisms accounted for decision making in our model, depending on the coherence level (Supply. Fig. S4). When plotting the expected winning onset rank of all association areas, we found a near-zero correlation (r=0.009) (Fig 3D). However, certain areas exhibited negative correlations in both scenarios (red dots in Fig. 3D), suggesting a reversal in sequential order upon task difficulty transitions. Specifically, their expected winning-onset rank showed a tendency to decrease along the hierarchy during low-coherence situations (top-down, r=-0.61), while increasing during high-coherence situations (bottom-up, r=0.65) (Fig. 3E and F, Suppl. Fig. S7A). Additionally, a small subset of regions remained stable across both scenarios (blue dots in Fig. 3D). We focused on the rank distribution of area MT to further illustrate the intrinsic mechanisms. Area MT experienced a remarkable transition from low to high winning-onset ranks when coherence increased from 0% to 100% (Fig. 3G). For low coherence, the differentiated sensory drive received by MT was weak, and due to the low self-coupling strength of this area, it took some time for its firing rate to reach the winning onset. Association areas, which typically have stronger self-coupling, reached their winning onset points faster in this condition, and hence ranking higher. For higher coherence levels, the differentiated sensory drive was strong enough to trigger a winning onset (and therefore a choice) in MT and related areas, and this information was then passed to higher association areas which followed MT.

The rank switch phenomenon could be further illustrated with a toy model (Fig. 3H, left) including only V1, MT and 9/46v. Here we rescaled the anatomical connections while keeping their relative ratios, and kept other parameters constant (see Methods). As in Fig. 2C, we observed activity ramping in both MT and 9/46v (Suppl. Fig. S5). Winning onset times decreased with coherence level for both areas (as easier trials lead to faster decisions), and the intersection between both curves signaled a switch in the winning onset ranking (Fig. 3H, right).

### Effects of receptor density per neuron and connectivity on evidence accumulation

The incorporation of NMDA and GABA receptor densities per neuron allowed us to explore their influnce on the evidence accumulation process. For the purpose of maintaining their linear relationship (Suppl. Fig. S1), we randomly shuffled NMDA and GABA receptor densities across the cortex as a whole. Following previous work (*33*), the maximum local excitatory self-coupling strength was set below the threshold value for a local bifurcation (=0.46). The network was then sensitive to certain parameters, particularly the global coupling G, which controlled the inter-area signal transmission strength (see Materials and Methods). When randomly shuffling parameter values, we set G to establish an adjusted network consistent with the controlled model, guided by two criteria: (i) before stimulus onset, all 40 areas remained in spontaneous states, and (ii) during the stimulus duration, with V1 excitatory populations receiving coherent directional inputs, the network exhibited ramping dynamics. After stimulus offset, some association areas displayed persistent activity indicative of the final decision.

We ran the simulation for 100 different parameter configurations and calculated the mean winning onset rank across 1000 trials. Interestingly, we found that the expected winning onset rank for each area displayed minimal difference under two task difficulty scenarios (Suppl. Fig. S6A). The correlations with the cortical hierarchy were -0.48 in low coherence situations and - 0.46 in high coherence situations, indicating a top-down modulation in both scenarios (Suppl. Fig. S6B and S7C). The computational mechanism behind this observation lies in the adaptive setup of individual excitatory background currents, which ensured that the spontaneous activities of each region remained constant. Consequently, external currents through inter-areal propagation did not change across varied parameter configurations. However, when high hierarchy areas like 9/46v, endowed with high excitatory self-coupling strength due to high dendritic spine count density, with their NMDA and GABA receptor densities shuffled with low hierarchy areas, the indirect inhibition effect between local excitatory population significantly decreased. This shift in the bifurcation point as a function of external current (Suppl. Fig. S6D) necessitated a decrease in global coupling strength to adjust the inter-areal current. As demonstrated in Suppl. Fig. S6E, the majority of fine-tuned global coupling strengths were significantly below the original model (G=0.52), indicating a less connected neocortical coupling. The loss of inter-areal coupling resulted in decreased efficiency of signal propagation. We recorded the long-range excitatory current in association areas in both scenarios under one shuffled receptor density setting and found that they were on the same scale as background noise (Suppl. Fig. S6F). Consequently, even in high coherence cases, the decision signal was insufficient to implement bottom-up transmission, and association areas would need to infer the final decision based on background noise and weak current decision signals, similar to low-coherence cases. We also applied the same protocol to shuffle the FLN connectome data and only the GABA receptor density data to disrupt the excitatory-inhibitory (E-I) balance. When shuffling FLN, the correlations between the expected winning onset and cortical hierarchy were nearly zero (Suppl. Fig. S7B). However, shuffling E-I ratio led to noticeable differences under both scenarios, with correlation coefficients of r = -0.59 under low-coherence cases and r = -0.23 under high-coherence cases (Suppl. Fig. S7D).

### Temporal gating in the macaque brain

In our large-scale cortical model, information about the decision taken can be stored in working memory as distributed sustained activity patterns, as in previous work (*33*) and experimental evidence (*42*–*44*). For categorical decision-making tasks, the robustness of such sustained activity is important for translating decision outcomes into motor output. This has been experimentally tested in rodents and macaques, by presenting distractors at different time points across the delay period, revealing that the robustness of decision outcomes greatly depends on the distractor timing, a phenomenon known as temporal gating (*45*–*47*). We tested the robustness of our decision-related sustained activity in a decision-making task with distractors (Fig. 4A), and here distractors were modeled as external currents into the V1 excitatory population different from the one encoding cue stimulus. Two versions of the task were considered: one in which distractors have a fixed onset but variable duration, and one in which they have a fixed duration but variable onset: at the sampling, early delay, or late delay phases (Fig. 4B).

**Figure 4:**
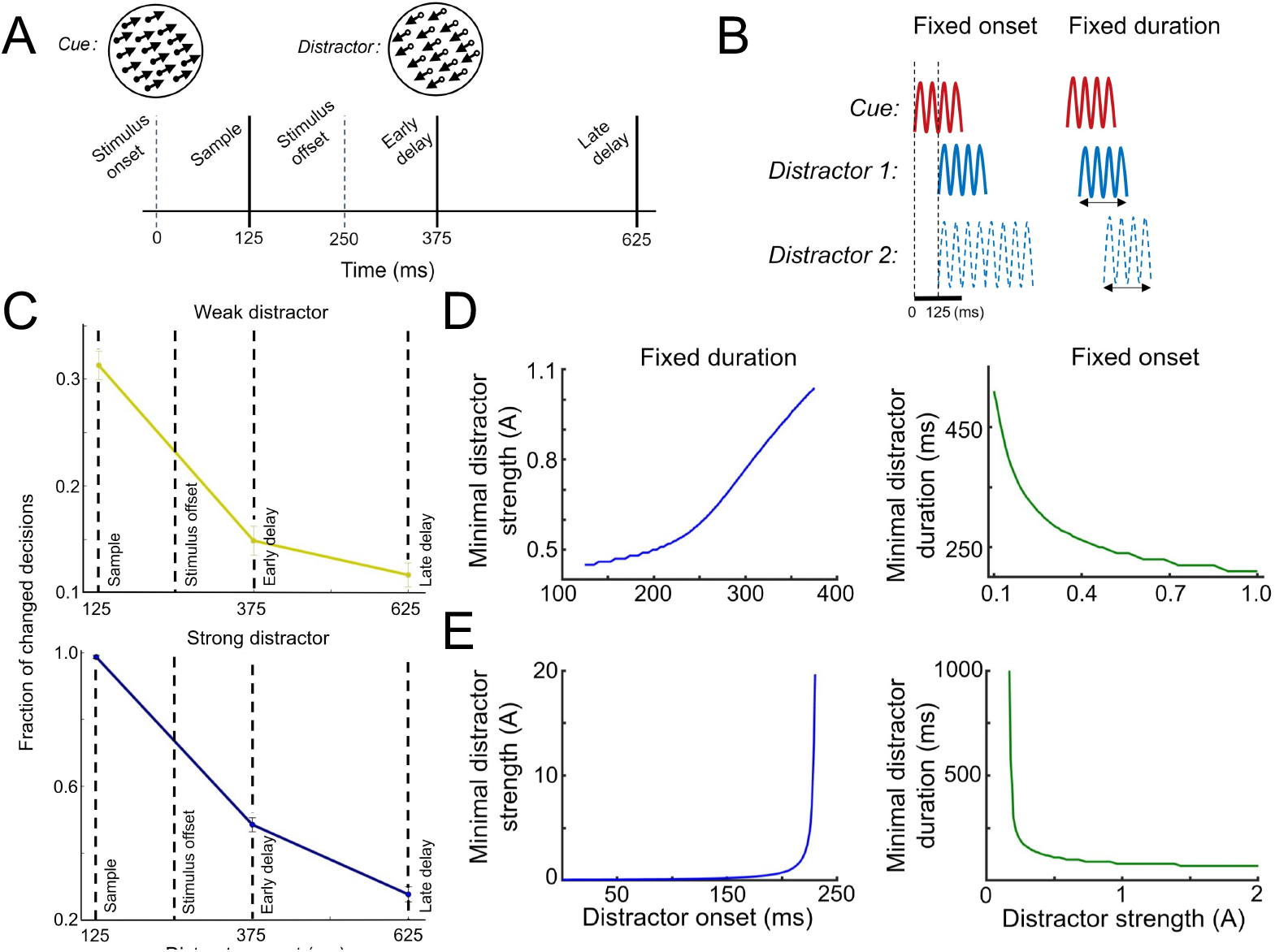
Temporal gating effects during decision making. (A) Decision making task with distractor. The model was presented with a 100%-coherence stimulus of strength 0.1 and duration 250ms, and then a distractor was presented in one of three time points (sample, early delay, or late delay). (B) Distractors in this task can be considered as either fixed onset (here at 125 ms) and variable duration, or as fixed duration (here, 250 ms) and variable onset. (C) Behavioral impact was quantified as the fraction of trials in which the network changed its decision due to the distractor, for the case of a weak (I=0.17, top panel) or strong distractor (I=0.80, bottom panel) and G=0.42. (D) Minimal effective distractors in terms of duration and strength, for the fixed duration (left) and fixed onset (right) versions of the task, global coupling G=0.42 and input noise of **σ**_**s**_ **= 0. 017**. (E) Same as panel D, but for a network with a stronger global coupling G=0.52 and **σ**_**s**_ **= 0. 01**.

We first studied the system in the fixed duration version of the task, and calculated the fraction of trials in which the decision was changed as a consequence of a distractor at different onsets (Fig. 4C). We observed that early distractors, especially those presented during the sampling phase, have stronger effects on changing the decision output than late ones, in agreement with experimental observations (*45, 46*). The effects were similar for weak and strong distractors, with the latter simply scaling up the fraction of trials with reversed decisions. We observed a positive relationship between the distractor onset and the minimal strength needed for the distractor to be effective, i.e. to change the decision outcome (Fig. 4D, left panel). For the fixed onset version of the task, we found an inverse relationship between the distractor strength and the minimal duration required for the distractor to be effective (Fig. 4D, right panel).

The results were similar for networks with a stronger global coupling (Fig. 4E), although with more abrupt limits in distractor effectiveness. For example, in the fixed duration version of the task (left panel), the decision was nearly impossible to be reversed with distractors of very late onset, irrespective of the distractor strength. Likewise, in the fixed onset version of the task (right panel), decisions could not be reversed with very weak distractors, irrespective of the distractor duration. Changes in distractor duration (for the fixed onset version) or distractor strength (for the fixed duration version) were determinant factors for the success of the distractor in overturning the decision (Suppl. Fig. S8).

### Inactivation reveals the functional irrelevance of LIP during decision making

Inactivation of specific brain regions, either by lesioning, pharmacology or optogenetics, has provided valuable insight for decision-making circuits. In macaques, inactivation via the GABA_A_ receptor agonist muscimol revealed a profound impact of MT inactivation in behavioral performance, while LIP inactivation had no measurable impact on performance(*21*). We used our model to simulate inactivation of MT and LIP areas (Fig. 5A), and we compared our modeling results with experimental evidence. We found a strong agreement between our results and the experimental findings of Katz and colleagues (*21*), with LIP inactivation barely affecting the psychometric curve for our model and MT inactivation leading to a flattening of the sigmoidal, reducing the accuracy of the decision process (Fig. 5B). Our results therefore support the evidence of LIP playing a relatively minor role in decision making, despite accurately reflecting the sensory accumulation process occurring across other distributed cortical networks. The effects of inactivation were consistent with (*21*) for higher values of the global synaptic coupling strength (G=0.52), but in this case, the effect of MT lesioning compared to baseline was less significant as the inactivation of any area could be easily covered by the joint contribution of other areas (Suppl. Fig. S9).

**Figure 5:**
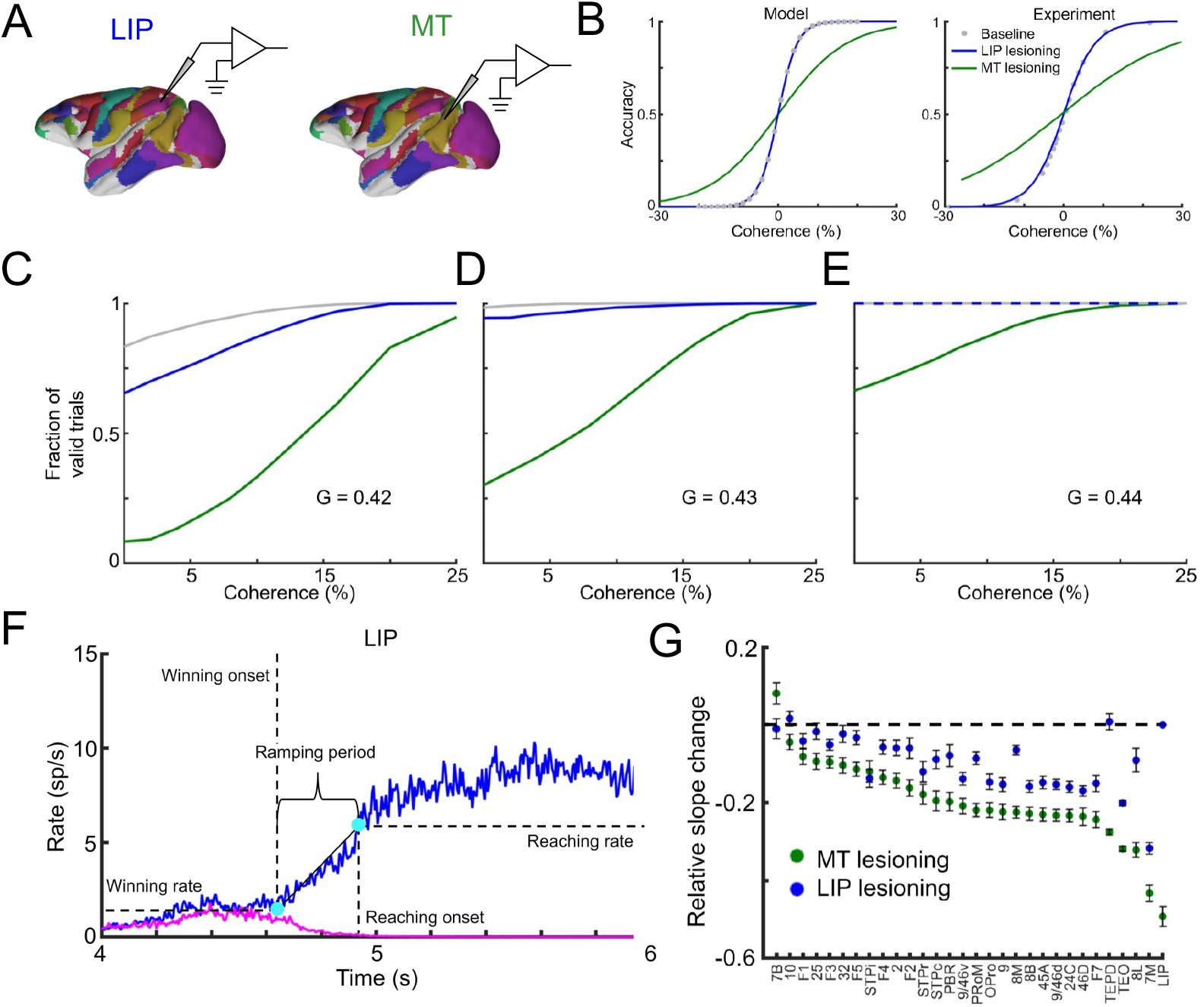
Comparison between MT lesion and LIP lesion in decision making. (A) Schematic of the inactivation protocol. (B) Psychometric curves for baseline, MT lesion and LIP lesion in macaques (left) and in our computational cortical model with global coupling strength of G= 0.42 and ***σ***_***s***_ **= 0. 018** (right). (C-E) Relationship between coherence and fraction of valid trials, for different global coupling strength values. (F) Illustration of winning onset, winning rate, reaching onset, reaching rate, and ramping period. (G) Relative slope change (percentage of ramping speed change) across individual areas when lesioning MT or LIP, averaged over 300 trials.

To better explain this result, we evaluated how the percentage of valid trials, defined as those for which the model was able to make a decision, depended on the coherence level for different global coupling values. For small global coupling, lesioning MT prevents the sensory evidence to reach higher cortical areas, leading to noise-driven trials and poor performance. Larger global coupling values ensure instead that trials are evidence-driven, leading to a good performance for MT inactivation due to compensatory effects of other regions. The inactivation of LIP, however, can be compensated by sensory accumulation in other areas for both small and large global coupling, leading to a causal irrelevance of LIP for the decision process (Fig. 5C-E). The role of MT as a bridge between sensory visual areas and superior association areas can be further elucidated by removing feedforward transmission in all 40 areas except from primary visual areas including V1, V2, V4 and V6, and then we injected a pulse input into the one of the excitatory populations in V1, resulting in responses exclusively from MT, TEO and TEpd to this stimulus (Suppl. Fig. S10). Considering the positions of MT, TEO and TEpd on the hierarchy axis were 8, 15 and 27, MT took a more important role in the transmission of visual information to higher areas.

Our distributed computational model was also able to infer how lesioning one area would affect the dynamics of other areas, especially their ramping speed, which is a key feature of the evidence accumulation process. To precisely define ramping speed, we introduced three area-specific metrics besides the winning onset: (i) the winning rate, defined as the firing rate of the winning population at the winning onset, (ii) the reaching rate, defined as 75% of the attractor firing rate minus 25% of the winning rate, and (iii) the reaching onset, defined as the time at which the winning population firing rate arrives at the reaching rate (Figure 5F). Ramping speed was defined as the change in firing rate over the ramping time window, and the lesioning effect of area *j* on area *i* was the percentage change between pre- and post-inactivation ramping speed. To better capture the ramping dynamics, we use a winning onset threshold of 0.5 instead of zero.

We then observed that lesioning MT had a stronger effect on other non-sensory areas than lesioning LIP (Figure 5G). Though lesioning LIP has a stronger impact in certain areas like 7B or STPi, the ramping speed of most areas (especially LIP, 7m, 8l and 10) changed more drastically when lesioning MT except for area 7B. Notably, lesioning LIP led to ramping speed increases in areas like F3 and TEpd; the reason was that long-range projections from LIP on those areas targeted inhibitory populations more strongly than excitatory ones, so removing LIP had a positive effect on those areas. This effect was not observed when lesioning MT, which follows from its early position on the feedforward cortical hierarchy and its role in the excitatory feedforward pathway.

### Effects of lesioning areas across the whole cortex

We extended the study of the previous section by lesioning all non-sensory areas one by one (Fig. 6A) and analyzed how any particular inactivation affects ramping speeds across cortex. The full inactivation matrix is displayed in Fig. 6C. We observed, for example, that lesioning prefrontal area 46d had a strong effect on the ramping speed of parietal areas LIP and 7m, indicating a supportive role of PFC on parietal cortex. Furthermore, lesioning TEO greatly decreased the ramping speed of TEpd and increased that of area 45A and the OPRO. Lesioning temporal areas like STPc, STPi and MT affected virtually all association areas.

**Figure 6:**
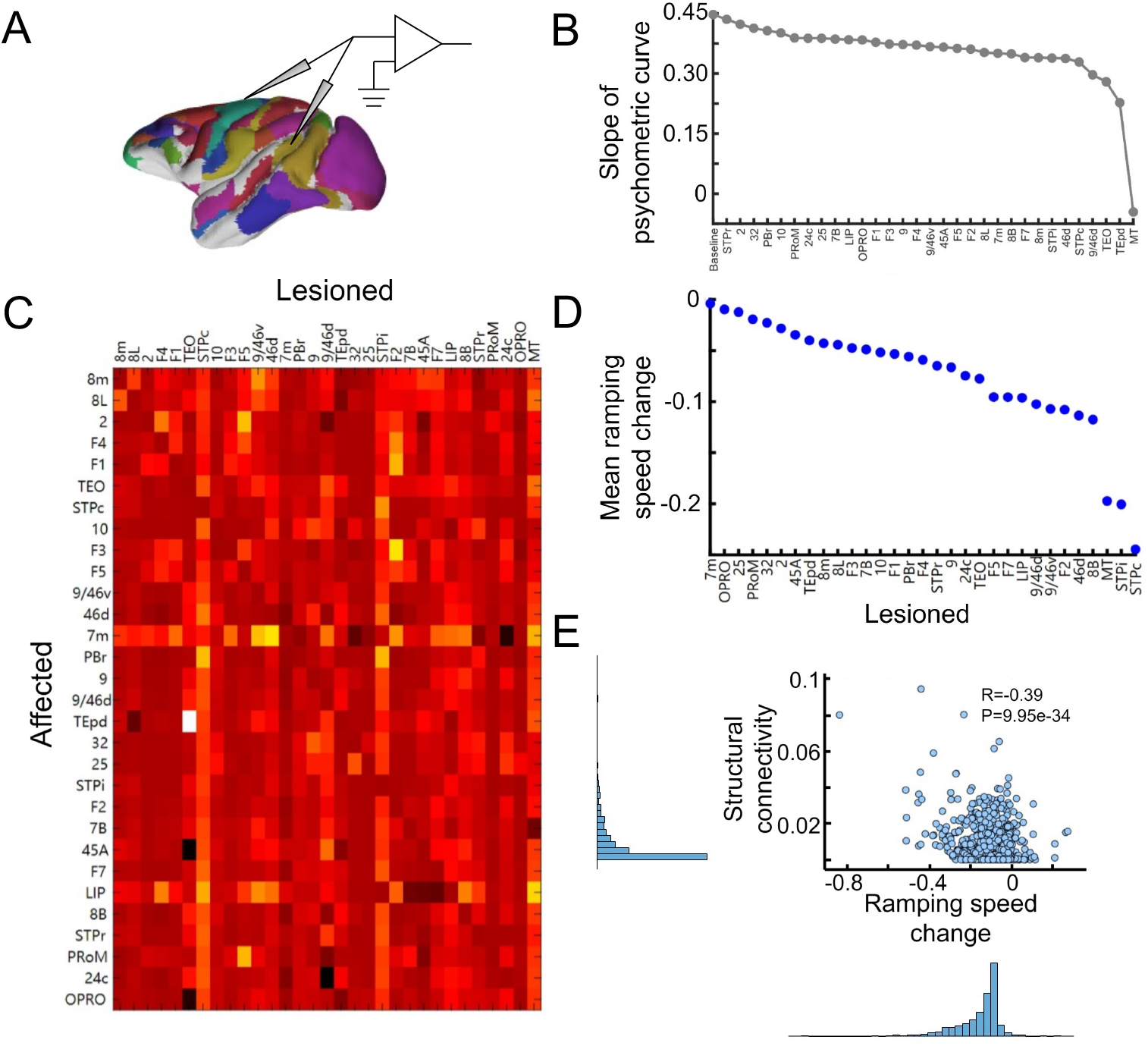
Model prediction of unrecorded cortical areas. (A) Schematic of the inactivation protocol for the neocortex. (B) Slopes of psychometric curve for baseline (0.45) and single area inactivation. (C) Inactivation matrix, with rows indicating the lesioned area, columns the affected areas, and color the relative slope change of the psychometric curve. (D) Mean of relative ramping speed changes when lesioning individual areas. (E) Relationship between lesioning effect (mean relative slope change) and structural connectivity. The distribution of corresponding variables is also displayed (left and bottom).

When we took mean values of the lesioning effect matrix, effectively applying the majority rule, we could infer how lesioning specific cortical areas globally affected the macaque’s decision-making performance (Fig. 6B) and the mean ramping speed across areas (Fig. 6D). For example, lesioning areas MT, TEO and TEpd had biggest effects on psychometric curve (as indicated by changes in its slope), corresponding to the ignition phenomenon when removing most feedforward transmission links (Suppl Fig. S8). Also, frontal areas like OPRO, PRoM, 32 and 25 had little effect on the psychometric curve, since their interareal connectivity with other regions was relatively weak. This link between structural connectivity (defined as *SLN* * *FLN*) and lesioning effects was reflected in a negative correlation of *r* = −0.39 between both factors (Fig. 6E). In contrast, inactivation of temporal areas led to strong effects, suggesting a more salient role of the temporal lobe in decision making.

### Discussion

In this work we investigated the distributed computations underlying decision-making tasks, by simulation of large-scale cerebral network models. Traditionally, decision making has been characterized by a gradual evidence accumulation process reflected in ramp-up activity in local cortical circuits (*6, 7, 48, 49*). While a few computational studies have approached this topic considering two interconnected areas (*27*–*29*), our work provides a formal leap as a novel and detailed computational study of decision making encompassing a large-scale, data-constrained cortical network. Additionally, our study opens the door to explore cognitive functions within large cortical networks in an integrative manner, as its architecture and properties align with a recent proposal regarding the distributed nature of working memory (*33*).

A salient feature of our work is the integration of multiple data sets of gradients of biophysical properties across the cortical network: (i) a pyramidal cell dendritic spine gradient informing on local synaptic density, (ii) a tract-tracing connectome embedding the model with directed and weighted long-range connections, (iii) a neuroanatomical cortical hierarchy used to infer targeted populations in long-range projections, and (iv) gradients of receptor density per neuron for the NMDA and GABA_A_ receptors which allowed to constrain the excitatory/inhibitory balance and strength. This level of detail allowed us to identify outliers like LIP or 7A, which had a high hierarchical rank and NMDA receptor density per neuron but a low level of pyramidal dendritic spines. This pointed at an unexpected abundance of NMDA receptors on inhibitory neurons, which contributed to the observed slow ramping activity in parietal cortex compared to other areas such as prefrontal cortex. The combination of data sets also allowed our model to explain sustained activity as reported in areas V4 and MT (*50, 51*), which previous models had missed. Despite this, our model, augmented with receptor heterogeneity, yielded different evidence accumulation trajectories compared to the hierarchy-driven model proposed by (*33*). For instance, the correlation between the expected winning onset rank of all association areas during low coherence trials exhibited only intermediate agreement between the two model simulations (Suppl. Fig. S11). Combining these data sets with additional refinements, such as causality-reconstructed connectivity maps and hierarchical interactions (*38, 52*–*55*) or additional receptor per neuron densities (*32, 39*) may further increase the model’s predictive capacity.

### Winning onsets uncover multiple information pathways

The sequential order of winning onsets observed in our model was a reflection of how motion information may be processed and broadcasted in the brain. Interestingly, the information pathways revealed by the model were not fixed, and rather dependent on the input coherence and therefore the task difficulty. We observed that higher cognitive and association areas like TEO and 9/46d led the decision process for hard trials, while early sensory areas led for easier trials. The alteration in winning onset order could indicate a change in the role of certain areas when the macaque encountered a tough vs simple task. This suggests that an efficient performance on difficult decision-making tasks might benefit from top-down modulation, a prediction aligned with conscious perception theories (*16*–*18, 56*). There is also evidence of monkeys taking different strategies depending on task difficulty (*57, 58*).

In addition, previous experimental attempts to uncover the information flow during categorical perceptual decision tasks revealed a decision-related cortical pathway given by MT→ LIP→ V4→ IT→ FEF→ PFC. By setting the input to full coherence, therefore approximating a categorical perceptual decision task, our model led to a similar cortical pathway for information flow attending to winning onset order: V4→ MT→ IT(TEO)→ LIP→ FEF (8I)→ PFC (9/46v) (Suppl. Fig. S3), with only minor misalignments with the data for the positions of V4 and TEO/IT. Furthermore, the activation of this sequence occurs within 0.2 seconds for both experimental data and our model predictions. This indicated that data-constrained large-scale models are well-suited to explore decision-related information pathways in cortical networks, and to uncover the mechanisms behind sequential decision stages (*45, 59*).

### Temporal gating in decision-related memory

In decision making tasks, monkeys are sometimes expected to maintain their choice in memory for a short time, which posits the robustness of choice-related memories as fundamental for performance. We varied global coupling strength to individually simulate both strongly and weakly distributed circuit models. As in the literature (*45*), distractors with different duration and strength were fed to the model, revealing distinct reactions to distractors and suggesting that early distractors are most effective than late ones. In weakly distributed models, it was easier for distractors to disturb the ramping activity, perhaps reflecting the performance of poorly trained monkeys. Future experiments could address how monkeys at different trained stages react to distractors, to test the existence of these two effects behind temporal gating (*60*).

### Causal irrelevance of LIP is replicated by simulated inactivation

Despite the traditional focus on parietal cortex as a circuit involved in sensory accumulation, recent evidence on rodents and primates has disputed this idea (*19, 21, 22, 25, 26*). However, a mechanistic explanation as of why parietal cortex would play a relatively minor role in decision making despite its activity’s high correlation with behavior was crucially lacking. By simulating lesioning effects across different areas, our model demonstrates that a distributed network of redundant evidence accumulation is sufficient to explain the experimentally observed causal irrelevance of LIP, as its inactivation may be compensated by the integration process occurring in parallel in other areas. Inactivation of other areas such as MT led by contrast to a much more substantial impairment in performance, as also observed experimentally. This is mostly due to the relatively low hierarchical position of MT, acting as an important bridge in the sensory pathway. When further exploring currently unknown effects of lesioning other areas, the model predicted an important role of temporal lobe areas such as TEO and TEpd, a prediction that requires experimental validation.

## Materials and Methods

The large-scale cortical model presented here uses previous modeling work on distributed working memory as a basis (*33*), and substantially improves upon it with the introduction of (i) a larger number of cortical areas interconnected by neuroanatomical data, (ii) area-specific data on the density of NMDA and GABA receptors across the neocortex, obtained via autoradiography experiments, and (iii) novel theoretical assumptions on the relative impact on the effects of the density of dendritic spines and postsynaptic receptors on intra- and inter-areal interactions.

We describe our model in three steps: the local circuit employed, the differentiation of such circuit across the whole neocortex via synaptic gradients, and the inter-areal projections. We then briefly define several metrics for behavioral performance and inactivation effects, ending with a short summary on our simplified ‘toy model’ for winning onset results.

### Computational model: local neural circuit

We employed the Wong-Wang model (*7*) to characterize the neural dynamics of a local microcircuit representing a cortical area. This model, in its three-variable version, captures the temporal evolution of the firing rates of two input-selective excittory populations, as well as the firing rate dynamics of an inhibitory population. The populations are interconnected with each other, as depicted in Figure 1A. The model is governed by the following equations:

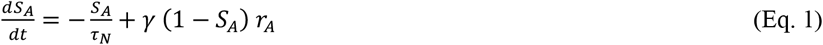

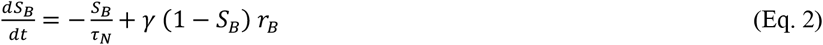

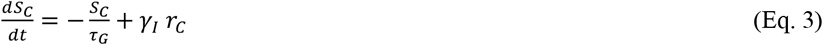

Here, S_A_ and S_B_ are the NMDA conductances of selective excitatory populations A and B respectively, and S_C_ is the GABAergic conductance of the inhibitory population. Values for the constants are τ_N_=60 ms, τ_G_=5 ms, γ=1.282 and γ_I_=2. The variables r_A_, r_B_ and r_C_ are the mean firing rates of the two excitatory and one inhibitory population, respectively.

They are obtained by solving, at each time step, the transcendental equation *r*_*i*_ = *ϕ*_*i*_(*I*_*i*_) (where *ϕ* is the transfer function of the population, detailed below), with I_i_ being the input to population ‘i’, given by

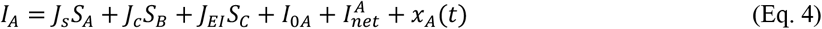

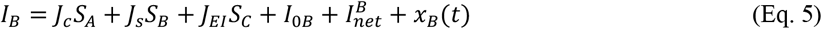

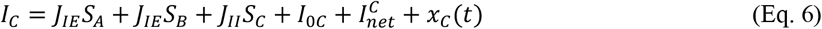

In these expressions, J_s_, J_c_ are the self- and cross-coupling between excitatory populations, respectively, J_EI_ is the coupling from the inhibitory populations to any of the excitatory ones, J_IE_ is the coupling from any of the excitatory populations to the inhibitory one, and J_II_ is the self-coupling strength of the inhibitory population. The parameters I_0i_ with i=A, B, C are background inputs to each population. Fixed parameters across the cortex are J_c_=0.0107 nA, J_EI_=-0.31 nA, J_II_=-0.20 nA, and I_0C_=0.26 nA (background currents for populations A and B are specified later). The term 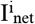 denotes the long-range input coming from other areas in the network and will be detailed later. Sensory stimulation can be introduced here as extra pulse currents of strength I_stimA_ =0.3(1+c) and I_stimB_=0.3(1-c) to the V1 excitatory populations A and B respectively, where c is the coherence level, with a duration of T_pulse_ =0.7 sec, unless specified otherwise.

The last term x_i_(t) with i=A, B, C is an Ornstein-Uhlenbeck process, which introduces some level of stochasticity in the system. It is given by

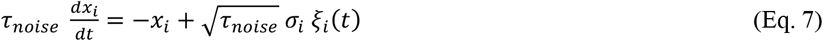

Here, ξ_i_(t) is a Gaussian white noise, the time constant is τ_noise_=2 ms and the noise strength is σ_A,B_=0.01 nA for excitatory populations and σ_C_=0 for the inhibitory one.

The transfer function ϕ_i_(t) which transform the input into firing rates takes the following form for the excitatory populations(*61*):

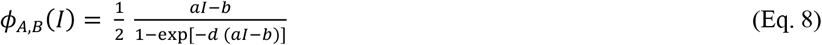

The values for the parameters are *a*=135 Hz/nA, *b*=54 Hz and *d*=0.308 s. For the inhibitory population a similar function can be used, but for convenience we choose a threshold-linear function:

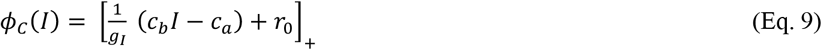

The notation [*x*]_+_ denotes rectification. The values for the parameters are g_I_=4, c_b_=615 Hz/nA, c_a_=177 Hz and r_0_=5.5 Hz.

### Computational model: Gradient of synaptic strengths

Before considering the large-scale network and the inter-areal connections, we look into the area-to-area heterogeneity to be included in the model.

Our large-scale cortical system consists of N=40 local cortical areas, for which inter-areal connectivity data is available. Each cortical area is described as a Wong-Wang model of three populations like the ones described in the previous section. Instead of assuming areas to be identical to each other, here we will consider some of the natural area-to-area heterogeneity that has been found in anatomical studies. For example, work from Elston (*31*) has identified a gradient of pyramidal cell dendritic spine density, from low spine numbers (∼600) found in early sensory areas to large spine counts (∼9000) found in higher cognitive areas. From an electrophysiological point of view, excitatory postsynaptic potentials (EPSP) have similar values both in early sensory (∼1.7±1.3 mV) and higher cognitive areas (∼0.55±0.43 mV). The combination of these findings suggests an increase of local recurrent strength as we move from sensory to association areas. In addition, cortical areas are distributed along an anatomical hierarchy (*37, 62*). The position of a given area ‘i’ within this hierarchy, namely h_i_, can be computed with a generalized linear model using data on the fraction of supragranular layer neurons (SLN) projecting to and from that area. In particular, we assigned hierarchical values to each area such that the difference in values predicts the SLN of a projection. Concretely, we assign a value H_i_ to each area A_i_ so that SLN(A_j_ → A_i_) ∼ f (H_i_-H_j_), with ‘f’ being a logistic regression. The final hierarchical values are then obtained by normalizing h_i_=H_i_/H_max_. Further details on the regression are provided elsewhere (*37, 38*).

In the following, we will assign the incoming synaptic strength (both local and long-range) of a given area as a linear function of the pyramidal dendritic spine count values observed in anatomical studies, with age-related corrections when necessary. Alternatively, when spine count data is not available for a given area, we will use its position in the anatomical hierarchy, which displays a high correlation with the spine count data, as a proxy for the latter. After this process, the large-scale network will display a gradient of local and long-range recurrent strengths, with sensory/association areas showing weak/strong local connectivity, respectively. We denote the local and long-range strength value of a given area *i* in this gradient as h_i_, and this value normalized between zero (bottom of the gradient, area V1) and one. We assume therefore a linear gradient of values of J_s_, with its value going from J_min_ to J_max_:

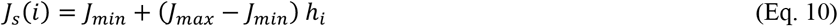

The above equation considers the impact of the number of dendritic spines per neuron in the synaptic strength of each brain area, but the density of NMDA and GABA_A_ receptors per neuron must also be taken into account. Since excitatory signals target not only other pyramidal neurons but also interneurons, we assume that the local NMDA receptor density has a positive correlation with the sum of all excitation signals in a given brain region. We use a saturating gradient, or logistic function, to model such area-specific parameters:

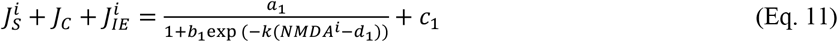

Here, the sum of the three excitatory projections in the left-hand side defines the strength of excitatory transmission, which depends on *NMDA*^*i*^, the experimental NMDA receptor density of brain area ‘i’, via a logistic function. The constants *a*_1_ = 0.524, *b*_1_ = 1.01, *c*_1_ = 0.018, *d*_1_ = 0.12, *k* = 9.35 are used to constrain the slope and roof of the curve. Since receptor density ratio between NMDA and GABA measures the relative excitatory-inhibitory synaptic strength balance, GABAergic receptor density can be introduced via the following balance equation:

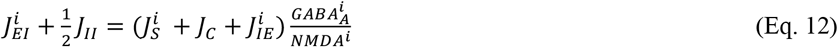

where *GABA*^*i*^ is the experimental GABA_A_ receptor density of brain area ‘i’.

If association areas have large values of *J*_*S*_, it can influence their spontaneous activity, even without considering inter-areal coupling. To ensure that the spontaneous firing rate of these areas remains within the physiologically realistic regime, a viable approach is to enforce that the fixed point of spontaneous activity is the same for all areas, which is a reasonable approximation. This can be done by adjusting the background currents *I*_0_ (for both excitatory populations A and B) on an area-specific basis, which aligns with the differentiated thalamocortical input observed in real brains. We therefore use adaptive background current to balance spontaneous firing rate:

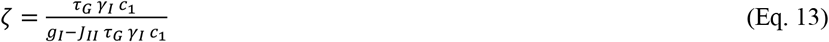

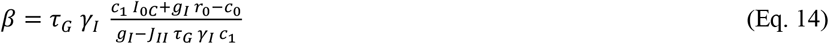

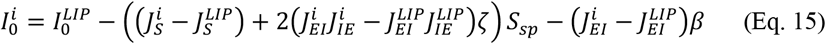

Here 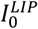 is the background current in parietal area LIP and its value has been set 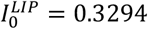 as in previous work (*7*) to fit temporal ramping excitatory neuron firing rates in LIP. The parameter *S*_*sp*_ = 0.03566 is the constant value of spontaneous NMDA gating variable in a local disconnected circuit.

Unless specified otherwise, we set *J*_*min*_=0.225 *nA* and *J*_*max*_ = 0.42 *nA* (i.e., below the critical value), so that the model displays distributed attractors as in the case of working memory (*33*).

### Computational model: Inter-areal projections

In the present study, the inter-areal projections that connect isolated areas contribute to the formation of the expansive cortical network. We assume that these inter-areal projections originate solely from excitatory neurons, as inhibitory projections typically exhibit a more localized pattern within real circuits and are selective for excitatory neurons. Thus, the network or long-range input term within a specific area “x” from all other cortical areas, can be expressed as follows:

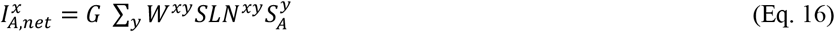

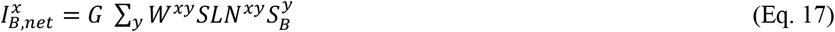

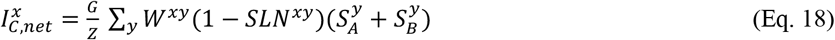

Here, *G* is the global coupling strength, which controls the overall long-range projection strength in the network (*G* = 0.52 unless specified otherwise). *Z* = 1.2 is a factor that considers the relative balance between long-range excitatory and inhibitory projections.

Aside from global scaling factors, the effect of long-range projections from population *y* to population *x* is influenced by two factors. The first one, *W*^*xy*^, is the anatomical projection strength as revealed by tract-tracing data (*36*). We use the fraction of labelled neurons (FLN) from population *y* to *x* to constrain our projections values to anatomical data. We rescale these strengths to translate the broad range of FLN values (over five orders of magnitude) to a range more suitable for our firing rate models. We use a rescaling that maintains the proportions between projection strengths, and therefore the anatomical information, that reads

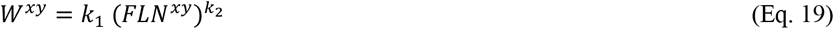

Here, the values of the rescaling are *k*_*1*_ =1.2 and *k*_*2*_ =0.3. The same qualitative behavior can be obtained from the model if other parameter values, or other rescaling functions, are used as long as the network is set into a standard working regime (i.e. signals propagate across areas, global synchronization is avoided, etc.) FLN values are also normalized so that ∑_*y*_ *FLN*^*xy*^ = 1. While in-degree heterogeneity might impact network dynamics (*63, 64*), this was done to have a better control of the heterogeneity levels of each area, and to minimize confounding factors such as the uncertainty on volume injections of tract tracing experiments and the influence of potential homeostatic mechanisms. In addition, and as done for the local connections, we introduce a gradient of long-range projection strengths using the spine count data: *W*^*xy*^ → (*J*_*s*_(*x*)/ *J*_*max*_) *W*^*xy*^, so that long-range projections display the same gradient as the local connectivity presented above.

The second factor that needs to be taken into account is the directionality of signal propagation across the hierarchy. Feedforward (FF) projections that are preferentially excitatory constitute a reasonable assumption which facilitate signal transmission from sensory to higher areas. On the other hand, having feedback (FB) projections with a preferential inhibitory nature contributes to the emergence of realistic distributed WM patterns (Figure 4) (see also previous work (*37, 65*)). This feature can be introduced, in a gradual manner, by linking the different inter-areal projections with the SLN data, which provides a proxy for the FF/FB nature of a projection (SLN=1 means purely FF, and SLN=0 means purely FB). In the model, we assume a linear dependence with SLN for projections to excitatory populations and with (1-SLN) for projections to inhibitory populations, as shown above.

Following recent evidence of frontal networks having primarily strong excitatory loops (*66*), it is convenient to ensure that the SLN-driven modulation of FB projections between frontal areas is not too large, so that interactions between these areas are never strongly inhibitory. In practice, such constraint is only necessary for projections from frontal areas to 8l and 8m (which are part of the frontal eye fields) and has little effect on the behavior of our model otherwise. The introduction of this limitation has two minor consequences: (i) it allows area 8l and 8m to exhibit a higher level of persistent activity during distributed WM –as their hierarchical position and recurrent strength are not strong enough to sustain activity otherwise, as previously suggested in anatomical studies (*36, 37*), and (ii) it slightly shifts the transition point in cortical space. Unless specified otherwise, we consider that the SLN-driven modulation of FB projections to 8l and 8m is never larger than 0.4.

### Behavioral performance: psychometric curves

Psychometric curves were fitted with a binomial generalized linear regression model (GLM):

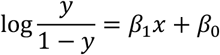

where *y* was the probability to choose population A as the final decision and *x* was the contrast strength. *β* = (*β*_1_, *β*_0_) were the parameters of the fit, with *β*_1_ representing the slope of the psychometric curve and *β*_0_ the value at zero contrast.

### Evaluation of lesioning effects

To replicate the results of the lesioning protocol, we employed a manual approach to induce lesioning effects in the target area’s three populations. Specifically, we set the firing rates of all three populations to zero. We evaluated the impact of such lesioning by assessing the percentage change in ramping speed, which measures the speed at which a specific area encodes the final decision:

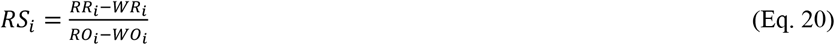

Here, RS_i_ is the ramping speed for area *i* in the decision-making task with the intact recurrent network. As mentioned in the main text, the winning onset (WO_i_) is the time at which the difference between the firing rates of both excitatory populations becomes larger than a certain threshold (zero for Fig. 3, 0.5 for Figs. 5 and 6) and increases from there without returning to zero (see Fig. 3B). The winning rate (WR_i_) is defined as the firing rate of the winning population at the winning onset, the reaching rate (RR_i_) is defined as (0.75 x attractor firing rate – 0.25 x winning rate), and the reaching onset (RO_i_) is defined as the time at which the winning population firing rate arrives at the reaching rate. We compare how lesioning one area affects the other area under same realization of Gaussian white noise with the following expression:

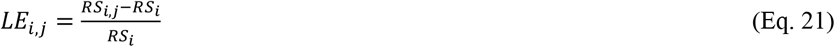

where LE_i,j_ is the degree by which lesioning area ‘j’ affects the ramping speed of area ‘i’, and RS_i,j_ is the ramping speed in area ‘i’ when area ‘j’ is lesioned.

### A toy model accounting for the shift of winning onset order

The toy model of Figure 3E consists of three cortical areas V1, MT and 9/46v, with V1 transmitting motion information, MT as an intermediate area connecting association areas with sensory areas, and 9/46v as an association area to sustain delay activity. The computational model structure is the same except for the long-range projections, which are now as follows:

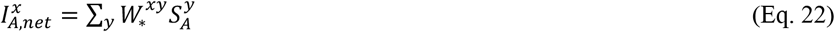

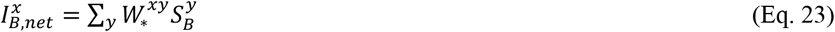

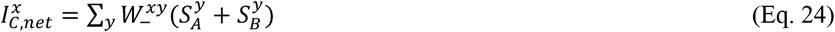

Here, 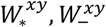 are parameters assigning individual roles to the three areas. All parameter values of the model can be found in Supplementary Table S1.

## Acknowledgments

We thank Kai Chen, Ziling Wang, Zhenyuan Jin, Chongming Liu for their support during the development of this work, Sean Froudist-Walsh for his help with data formatting and visualization, and Henry Kennedy for providing the connectivity dataset.

## Author contributions

JFM designed the study; NPG contributed datasets; LZ performed the research; LZ, DZ, SL and JFM analyzed and discussed the results; LZ, NPG, DZ, SL and JFM wrote the manuscript.

## Data and materials availability

All information needed to reproduce the results of this manuscript are in the main text and Materials and Methods section, and the code used will be made available upon publication of this work.

## Supplementary figures

**Figure S1:**
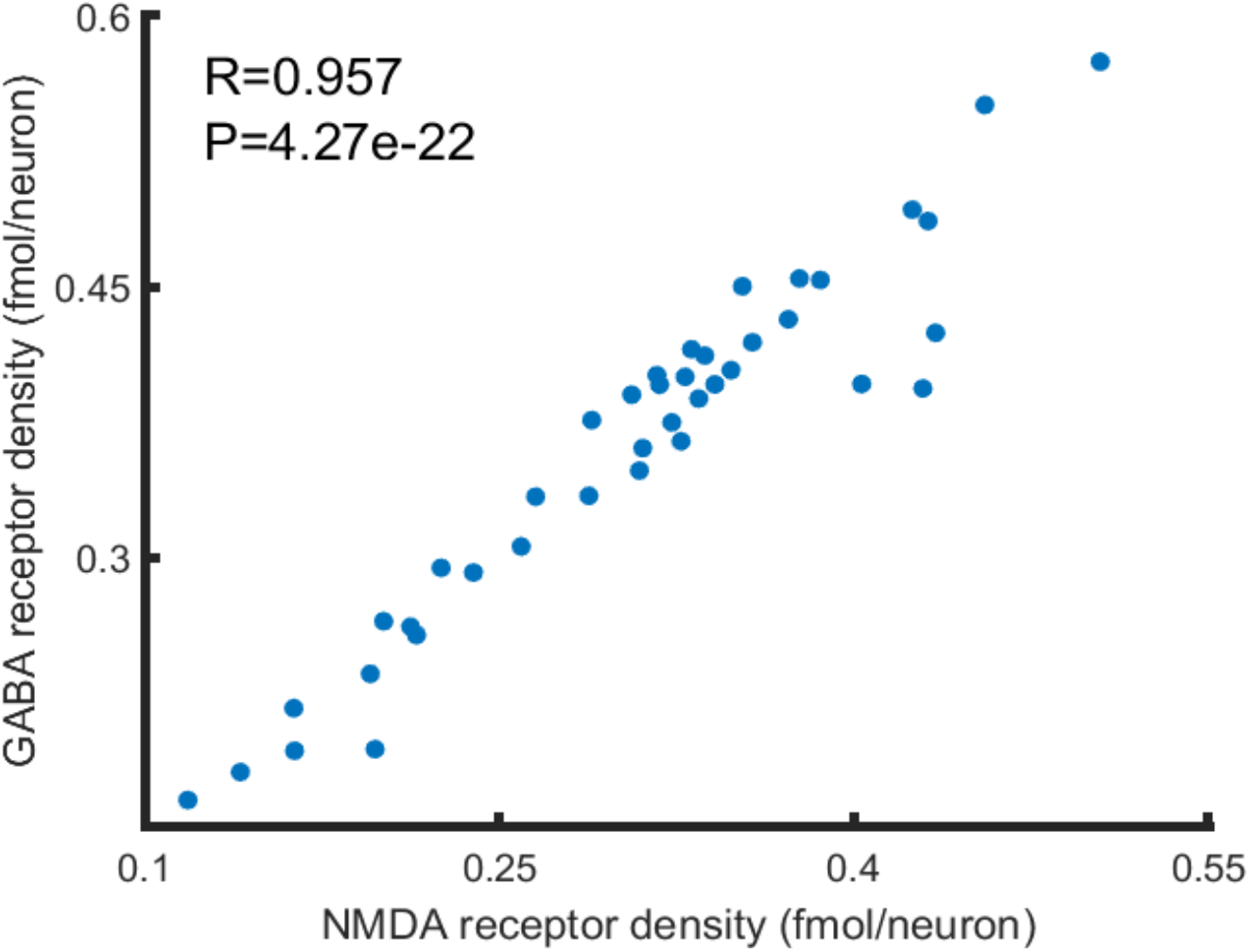
Correlation between the density of GABA_A_ and NMDA receptors per neuron across all cortical areas considered. The high correlation allows to adopt several modeling assumptions (see Methods).

**Figure S2:**
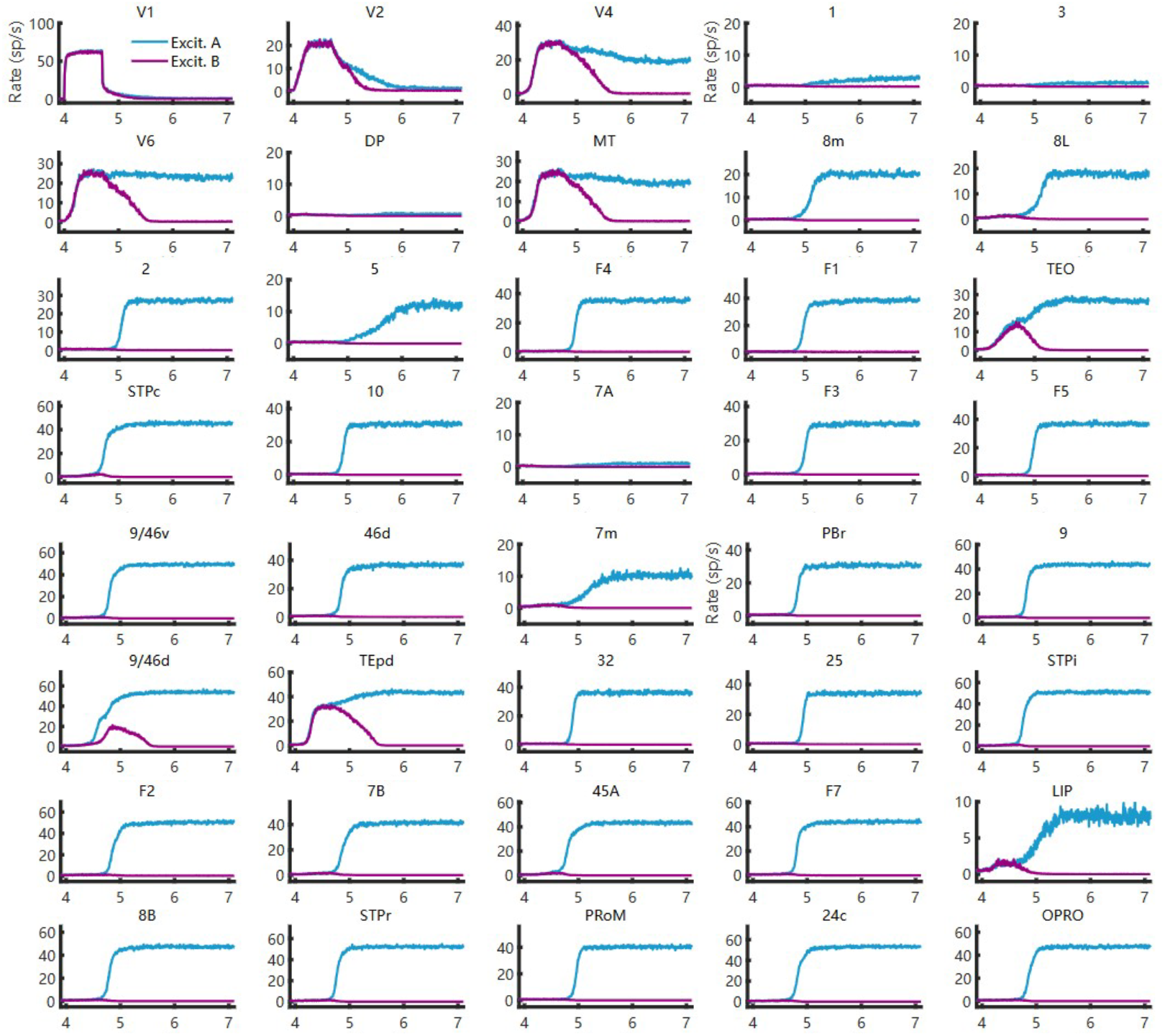
Evidence accumulation is distributed across multiple cortical areas. Integration of sensory evidence across all 40 cortical areas in the model, for a visual input of 5% coherence entering V1. We can distinguish between sensory-driven areas (such as V1 and V2), accumulators (like TEO, LIP and 9/46d) and classificators (like 45A and 24c) depending on their response profile and speed of integration.

**Figure S3:**
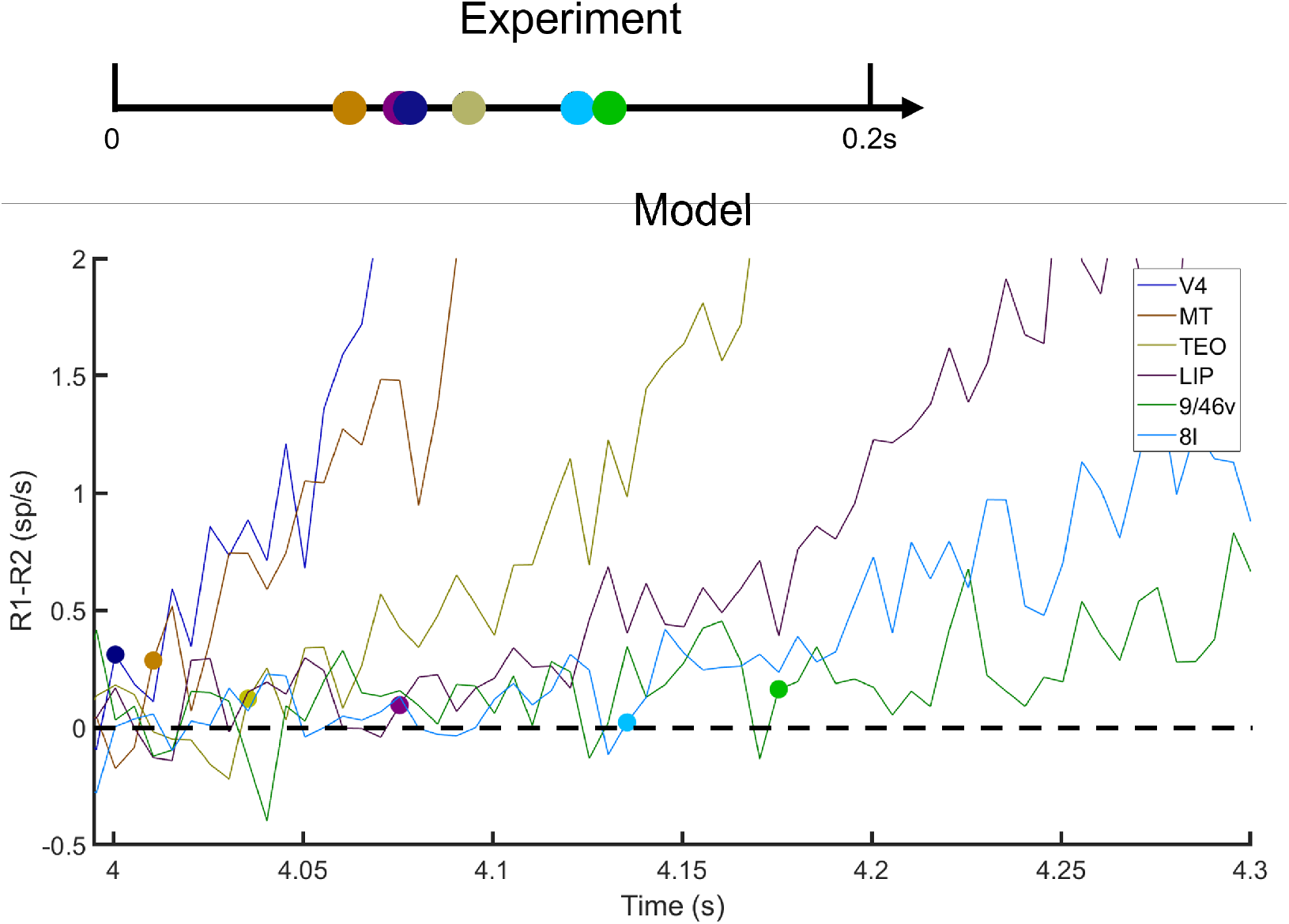
Comparison of area-specific decision times in experiments and model. Top: Timing of onsets in the cortical information flow in categorical decision-making as measured experimentally in macaques **(*9*)**. Bottom: Timing of the winning onsets for decision making as predicted by our model. Area legend applies to both panels. Both experimental data and model predictions have the same overall activation window of ∼0.2 seconds and both follow the same order, except for areas V4 and TEO/IT.

**Figure S4:**
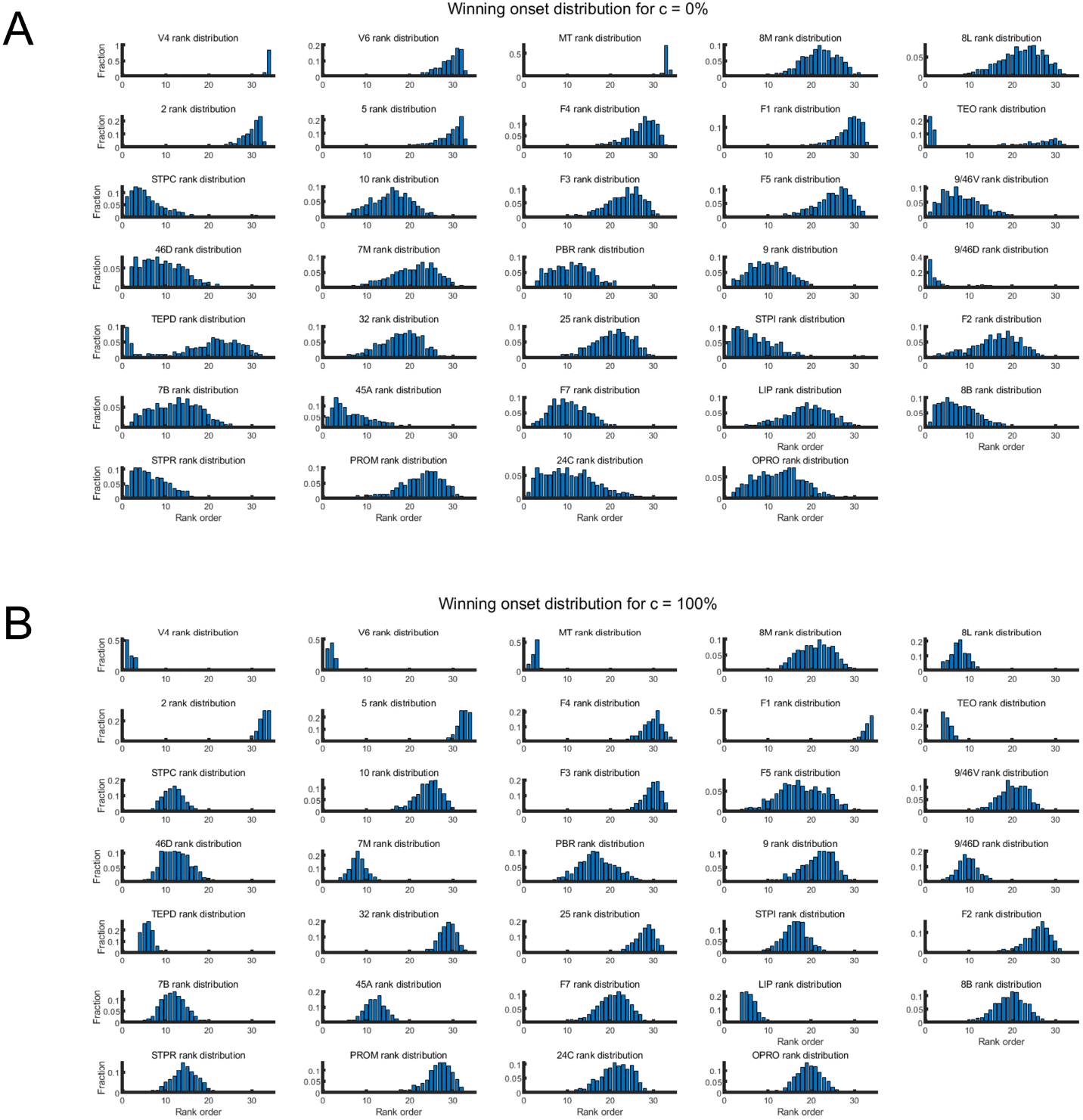
Winning onset distribution of all association areas: (A) Coherence = 0%. (B) Coherence = 100%.

**Figure S5:**
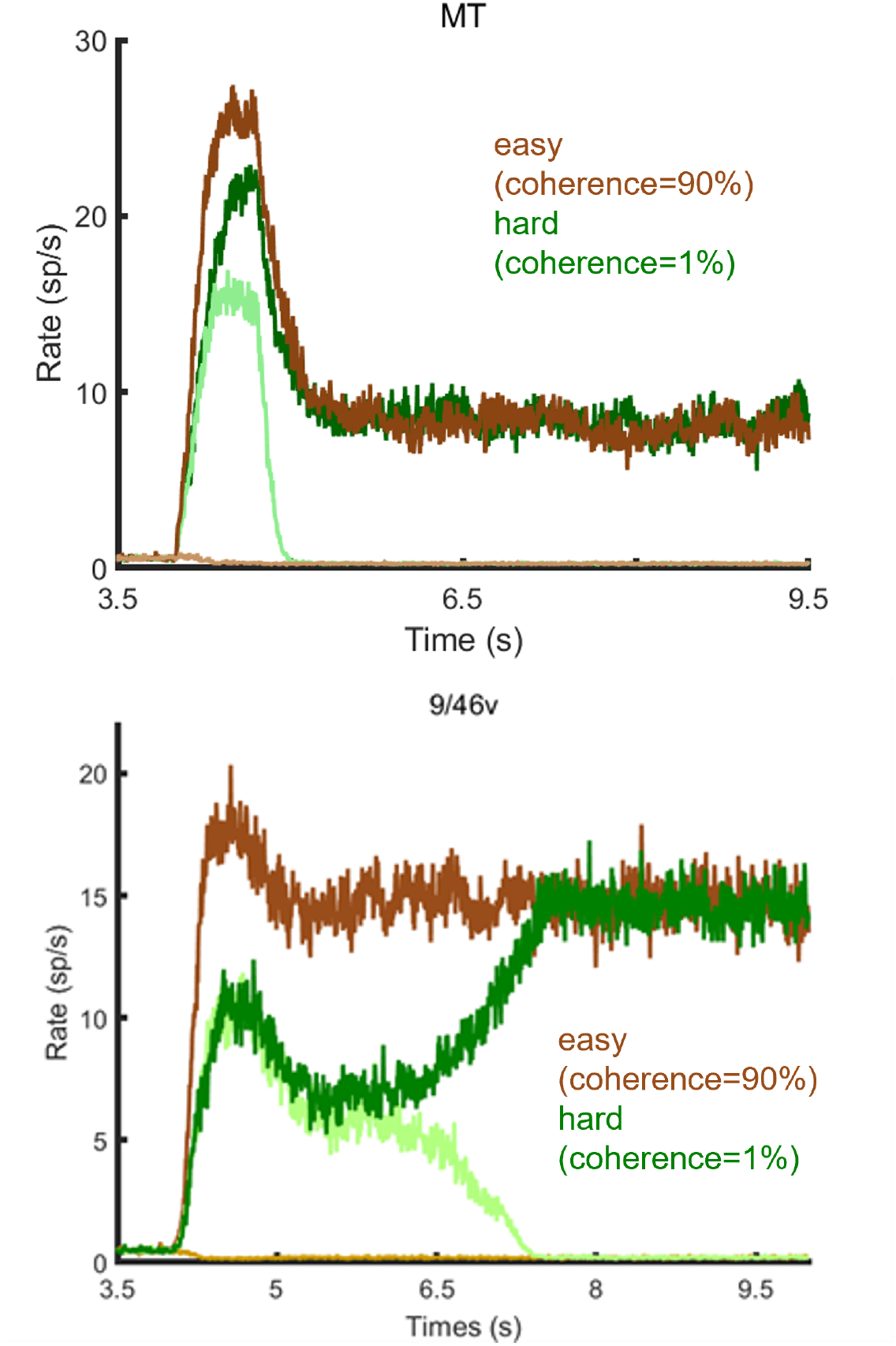
Evidence accumulation at MT and 9/46v during the decision-making task for the simplified three-area model (V1, MT, 9/46v). As in the full-network model, trials with low coherence led to longer decision times.

**Figure S6:**
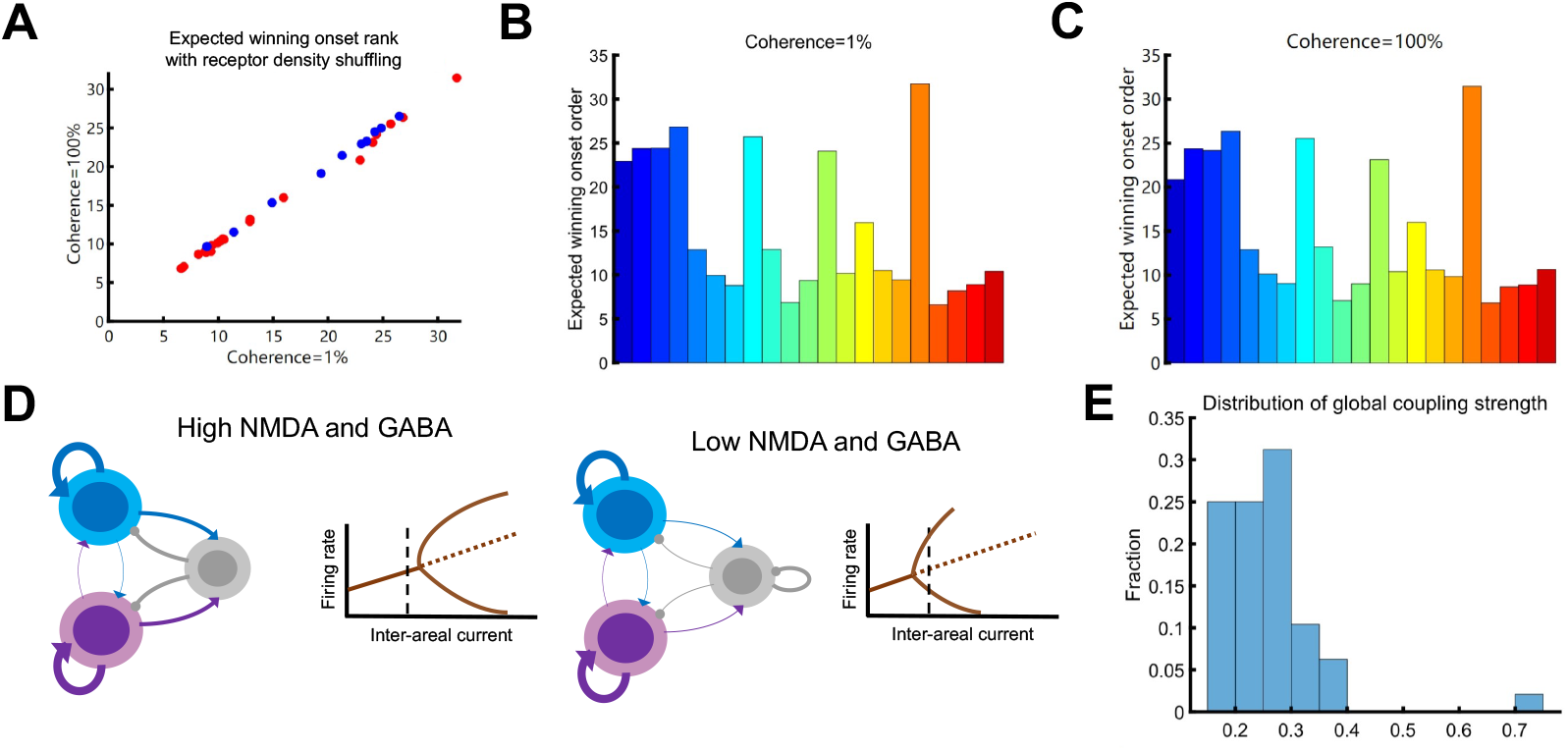
Effects of NMDA and GABA receptor density shuffling. (A) The expected winning onset ranks under coherence 1% and coherence 100% of 32 association areas, with V4 and MT excluded. (B) Relationships between expected winning onset ranks and cortical hierarchy position under coherence 1%. (C) The same as (B), under coherence 100%. (D) The bifurcation diagram for a specific area with high dendritic spine count, high NMDA and GABA receptor densities. In the control model (left), the inter-areal current is not enough to induce bifurcation at spontaneous states. When shuffling its receptor density data with a lower area, with the same global coupling strength, the inter-areal current crosses the new bifurcation point (right) because of the decreasing indirect inhibitory coupling between excitatory populations. Dashed line is the spontaneous inter-areal current. (E) Distribution of global coupling strength over 100 receptor density data shuffling. (F) For one specific shuffled model configuration, the neocortical responses for stimulus under coherence 1% (left) and coherence 100% (right). (G) The corresponding dynamics of inter-areal current in both scenarios. (H) Temporal dynamics of background excitatory current noise, on the same scale compared to inter-areal current in (G).

**Figure S7:**
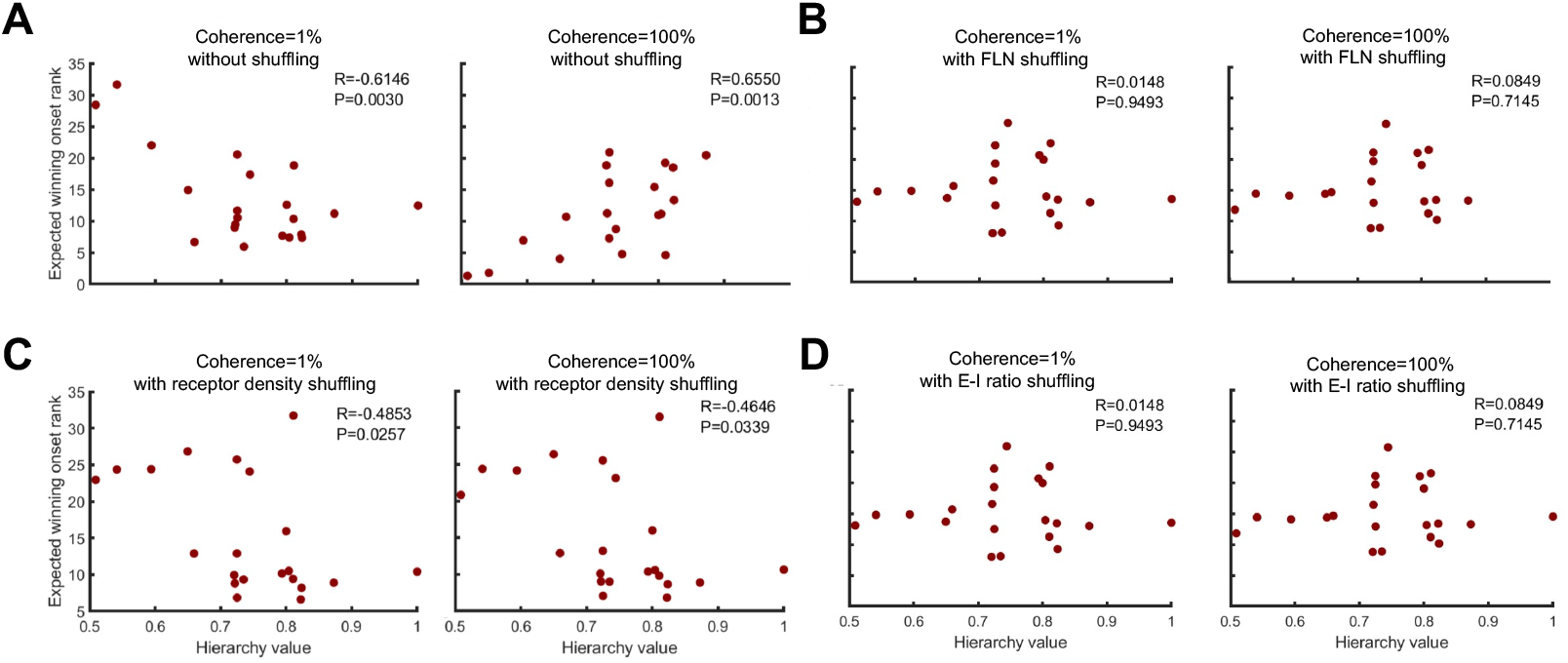
Correlation between expected winning rank and hierarchy value under different shuffling situations. (A) Without shuffling. (B) Shuffling FLN connectome. (C) Shuffling NMDA and GABA receptor density together. (D) Shuffling E-I ratio (only GABA receptor density).

**Figure S8:**
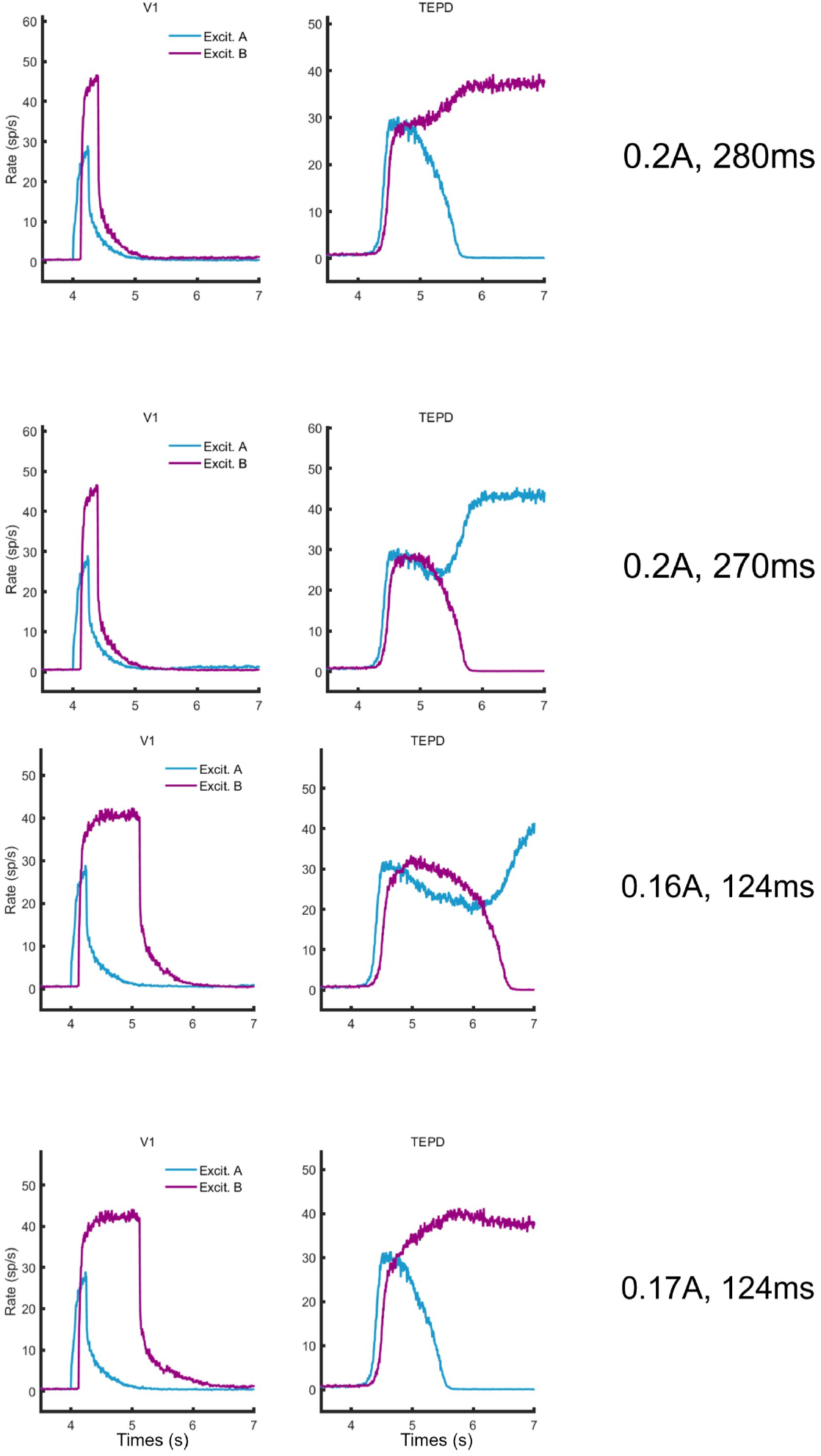
Temporal gating effects. Impact of temporal gating of distractor signals (purple) on the maintenance of cue information (blue) for V1 and TEpd. Top and middle-top: effects of changing the duration from 280 ms to 270 ms in a fixed onset task. Middle-bottom and bottom: effects of changing the distractor strength in a fixed duration task.

**Figure S9:**
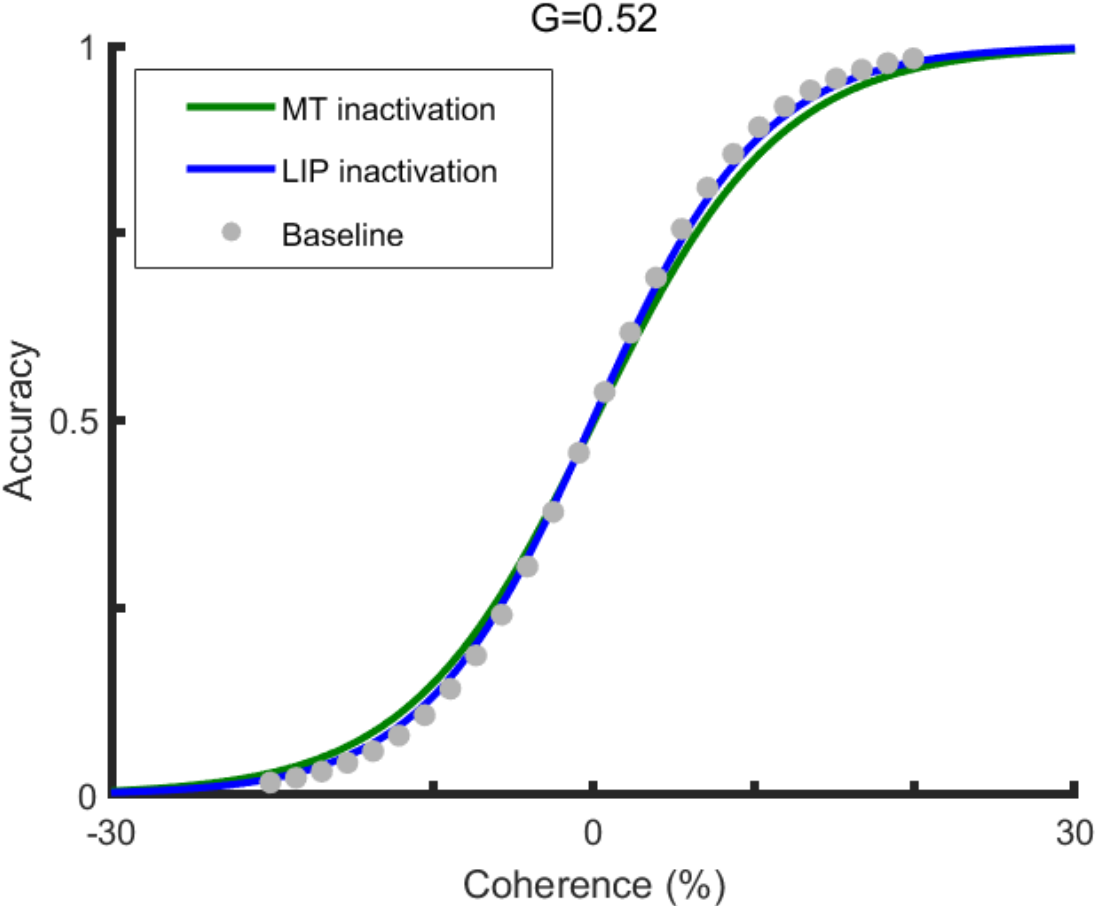
Lesioning effect on behavioral performance for strongly connected networks. Psychometric curve of the full-network model for the control case (baseline), inactivation of MT (green) and inactivation of LIP (blue) for global coupling of G=0.52. Due to the strong interaction between cortical areas, lesioning either MT or LIP does not lead to substantial performance drops in the task, in contrast to the case of weakly connected networks described in the main text.

**Figure S10:**
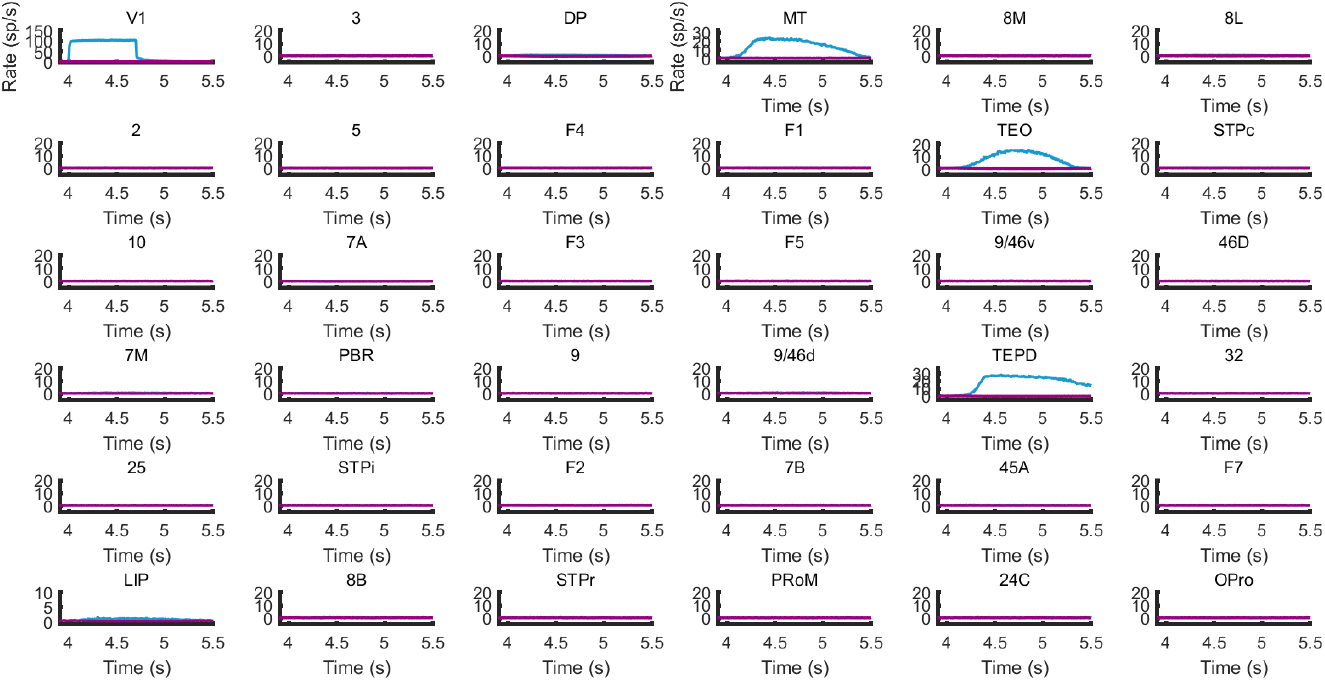
Signal propagation of visual information. Feedforward connections were all removed except originating from visual areas V1, V2, V4 and V6, and a step visual input (=0.6A) was injected into one of V1 excitatory populations. All association areas didn’t respond to this stimuli despite MT, TEO and TEpd, indicating the role of visual transmission bridges of those areas.

## Supplementary tables

**Table S1.** Parameters of the toy model used to explain the shift in winning onset values.

|  | **V1** | **MT** | **PFC** |
| --- | --- | --- | --- |
| **J<sub>s</sub>** | 0.25 | 0.42 | 0.29 |
| **J<sub>IE</sub>** | 0.015 | 0.05 | 0.1 |
| **I<sub>0I</sub>** | 0.26 | 0.26 | 0.26 |
| **I<sub>0E</sub>** | 0.3195 | 0.3192 | 0.3172 |
| **W<sub>E</sub> from V1 to area:** | 0 | 0.07 | 0.01 |
| **W<sub>E</sub> from MT to area:** | 0.01 | 0 | 0.1 |
| **W<sub>E</sub> from PFC to area:** | 0.01 | 0.07 | 0 |
| **W<sub>I</sub> from V1 to area:** | 0 | 0.001 | 0.01 |
| **W<sub>I</sub> from MT to area:** | 0.01 | 0 | 0.005 |
| **W<sub>I</sub> from PFC to area:** | 0.01 | 0.05 | 0 |

